# Spatiotemporal Analysis Reveals Mechanisms Controlling Reactive Oxygen Species and Calcium Interplay Following Root Compression

**DOI:** 10.1101/2025.10.22.683952

**Authors:** Pauline Vinet, Vassanti Audemar, Pauline Durand-Smet, Jean-Marie Frachisse, Sébastien Thomine

## Abstract

Mechanical stimulation of the root triggers signal transduction involving Reactive Oxygen Species (ROS) and calcium, but their relationships are unclear. This study aims to clarify the temporal and spatial interrelations between calcium and ROS following a localized lateral compression of the root. We combined a microfluidic valve rootchip to apply controlled compression, with fluorescent probes and wide-field or confocal microscopy to monitor H_2_O_2_ and calcium dynamics in root tissues simultaneously. Pharmacological inhibitors were used to investigate the causal links between H_2_O_2_ and calcium responses. In response to compression, we observed transient H_2_O_2_ accumulation, with characteristics similar to the calcium response observed previously in the same microfluidic system. H_2_O_2_ and calcium response occurred in 3 kinetic phases: a fast calcium increase relying on mechanosensitive channels and external calcium entry, followed by a long-lasting H_2_O_2_ accumulation and a slow calcium increase depending on NADPH oxidase activity. H_2_O_2_ accumulated in all root tissues while calcium increases were confined to the root center. These results suggest that two mechanotransduction mechanisms are involved in root response to compression. One mechanism relies on plasma membrane mechanosensitive channels, triggering a fast calcium increase. Another independent mechanism, relying on FERONIA, induces H_2_O_2_ accumulation, which drives the slower secondary calcium increase.

## Introduction

Plants are frequently exposed to mechanical stimuli, such as touching of the leaves or bending of the aerial parts caused by the wind. Roots also experience mechanical stress when growing against an obstacle or when compressed laterally by soil compaction or upon bending (Monshausen et al., 2009; Kolb et al., 2012). In order to adapt to their mechanical environment, plants sense and transduce mechanical stimuli into biological responses. Long-term responses mediated by changes in gene expression, thigmomorphogenesis or lateral root organogenesis can be distinguished from rapid/short-term responses such as ROS production, intracellular calcium elevations and pH changes (Braam and Davis, 1990; Monshausen et al., 2009; Richter et al., 2009).

Intracellular calcium (Ca²⁺) and ROS are ubiquitous mediators of rapid plant responses to biotic and abiotic stresses (Gao et al., 2004; Ranf et al., 2011; Ravi et al., 2023). In response to abiotic stress, ROS and calcium waves propagate in the roots and shoots, allowing systemic signaling (Evans et al., 2016; Fichman et al., 2022). Calcium responses in various organelles display specific kinetics, forming unique calcium signatures. ROS responses involve different types of ROS (O_2_**^.-^**, ^1^O_2_, H_2_O_2_, HO.) that accumulate in different cell compartments (Mittler et al., 2022). In the root, monophasic Ca²⁺ dynamics occur in response to touch, while the response to bending is biphasic (Monshausen et al., 2009). In addition, side-specific signatures have been reported in roots upon bending (Shih et al., 2014). Similarly, in the leaf epidermis, pavement cells can distinguish between mechanical touch and release, exhibiting distinct calcium signatures depending on the type of mechanical stimulus (Howell et al., 2023). Altogether, this points to stimulus-dependent regulation of the calcium response.

The link between ROS and calcium responses has been previously characterized in roots under salt stress and in leaves in response to light, pathogen, salt, or wounding (Ranf et al., 2011; Evans et al., 2016; Fichman et al., 2022; Marcec and Tanaka, 2022). The interplay between ROS and Ca²⁺ varies according to stress type: in flagellin 22–treated leaves, each signal modulates the other (Marcec and Tanaka, 2022); in response to microbe- or damage-associated molecular patterns, ROS influence calcium signatures (Ranf et al., 2011); and under salt stress, ROS and Ca²⁺ waves mutually amplify (Evans et al., 2016).

In response to mechanical stimulation, ROS and Ca²⁺ responses are triggered within the same timescale (Monshausen et al., 2009; Shih et al., 2014; Kulich et al., 2026). In addition, the mechanosensitive channel MSL10 is required for calcium and ROS responses induced by cell swelling (Basu and Haswell, 2020). Together, these previous studies highlight the interconnected responses of ROS and Ca²⁺ to mechanical stimulation. In a more detailed investigation, Monshausen *et al*. showed that ROS production in response to touch is abolished by inhibition of calcium channels, and that calcium entry is sufficient to induce a ROS response, showing that calcium triggers ROS production (Monshausen et al., 2009). However, whether and how ROS affect the calcium response has not been investigated, and the spatial and temporal interdependence of Ca²⁺ and ROS responses to mechanical perturbation remains to be clarified.

To investigate cell signaling in response to root compression, we combined mechanical compression of a root with a microfluidic valve with real-time monitoring of ROS and cytoplasmic Ca²⁺. The microfluidic valve rootchip allows controlling the intensity and duration of the local mechanical stimulation applied to the root with a high temporal resolution. This device has been used to investigate calcium signatures in response to compression (Audemar et al., 2023). Here, we first show that compression of the root triggers a local and transient ROS accumulation whose amplitude decreases upon repeated stimulations. The ROS response exhibits similarities with calcium signatures observed previously using the same experimental setup (Audemar et al., 2023). High temporal resolution monitoring of ROS and calcium in the same root revealed three phases in the response: first, a fast increase of calcium lasting a few seconds, then a slower ROS accumulation and finally an even slower calcium increase. The ROS response was ubiquitous across all root tissues, while both calcium phases localized specifically in the root center. The fast calcium peak required mechanosensitive channels and the entry of external calcium. Conversely, the sustained ROS wave and the subsequent slow calcium phase were partially dependent on ROS production generated by NADPH oxidases. These two phases were also partially dependent on FERONIA.

## Results

### The ROS response to root compression is similar to the previously observed calcium response

Audemar et al. previously reported calcium increase in response to a lateral compression of the root with a microfluidic valve rootchip (Audemar et al., 2023). We wondered whether the cells also react with the accumulation of ROS, which is often associated with calcium increase in mechanotransduction. H_2_O_2_ production was visualized using 10 µM Amplex Red probe added in the medium flow in root channels (Fig. 1A). The Amplex Red signal observed had both intracellular and extracellular localizations in our experiments (Supplementary Fig. 2). Amplex Red intensity in the root region of interest (Fig. 1B) was measured in response to compressions triggered by the application of 90 kPa pressure in the channels overlayed on the root (Fig. 1A, Supplementary Fig. 1). Compression was always applied in the mature zone of the root. This pressure level induces a 10 % deformation in root diameter along the axis of pressure application (Audemar et al., 2023).

**Figure 1:**
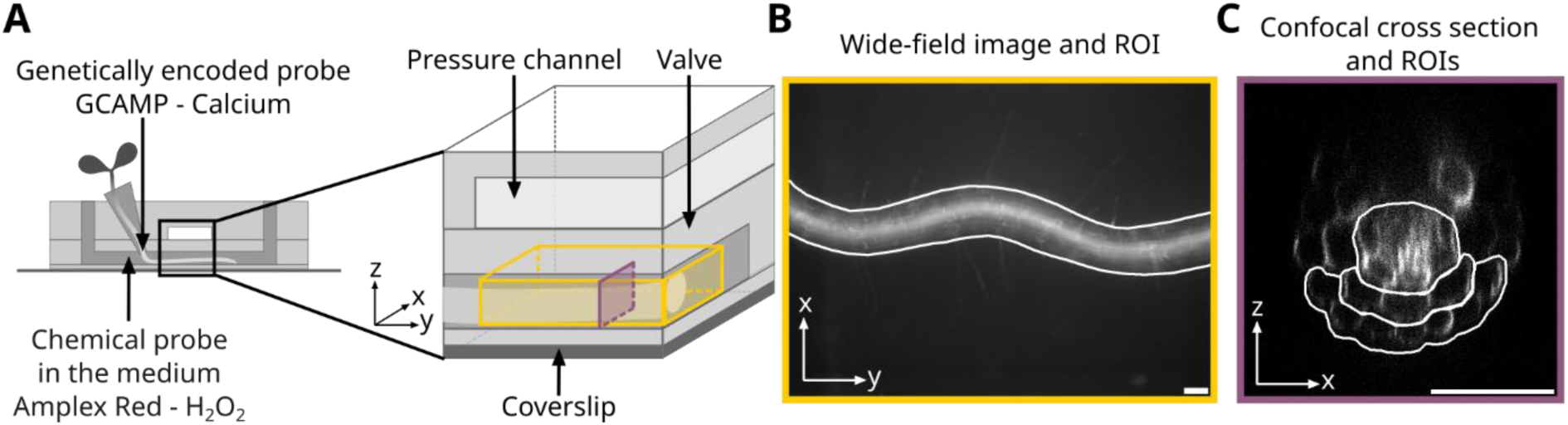
Microfluidic valve rootchip and imaging techniques to study H_2_O_2_ and Calcium response to a lateral compression. **(A)** 2D view and 3D zoom on the microfluidic device. The valve is a thin deformable layer separating the pressure channel and the root channel. Yellow volume represented on the 3D zoom corresponds to the volume imaged for the wide-field microscopy experiments at 10X magnification. The purple plane corresponds to the 20X magnification confocal cross-sections imaged during the tissue localization experiments. Roots were loaded with 10µM Amplex Red for H_2_O_2_ measurements and seedlings expressing GCAMP3 were used for calcium measurements. **(B)** Example of an XY image of a root loaded with Amplex Red obtained in wide-field microscopy. The white outlines indicates the region of interest where fluorescence intensity is measured. Scale bar 50 µm. **(C)** Example of an XZ confocal cross-section of a root loaded with Amplex Red. White outlines indicate regions of interest corresponding to the epidermis, cortex and root center zones for which fluorescence intensity was quantified. Scale bar 50 µm.

In response to a short 1-minute-long compression, we observed a rise in Amplex Red fluorescence in the root zone under the pressure valve, which did not propagate away from the stimulated area (Fig. 2A-D, Supplementary Fig. 1C). This rise in Amplex Red signal indicated local H_2_O_2_ accumulation in response to lateral compression of the root. This accumulation was transient, reaching a maximum 61 ± 4 s after pressure start. The mean fluorescence intensity returned to the initial value within about 5 minutes (Fig. 2C, for the first compression). Amplex Red probe is irreversibly converted into a fluorescent residue (resorufin) in presence of H_2_O_2_ and peroxidases. The reversibility of the signal observed in the microfluidic system is due to washout of the resorufin by the continuous flow of fresh medium in the root channels, as stopping the medium flow led to a steady increase in fluorescence, which decreased when perfusion was resumed (Supplementary Fig. 3). Our system thus measures the balance between Amplex Red oxydation to resorufin by peroxidases and fast resorufin washing out by the perfusion at the constant rate imposed by syringe pumps. This may lead to underestimation of the kinetics of H_2_O_2_ production but not the onset time of this response. These results show that root lateral compression induces a transient increase in ROS accumulation that does not propagate outside of the stimulated zone, with similar kinetics as the calcium response already observed by (Audemar et al., 2023).

**Figure 2:**
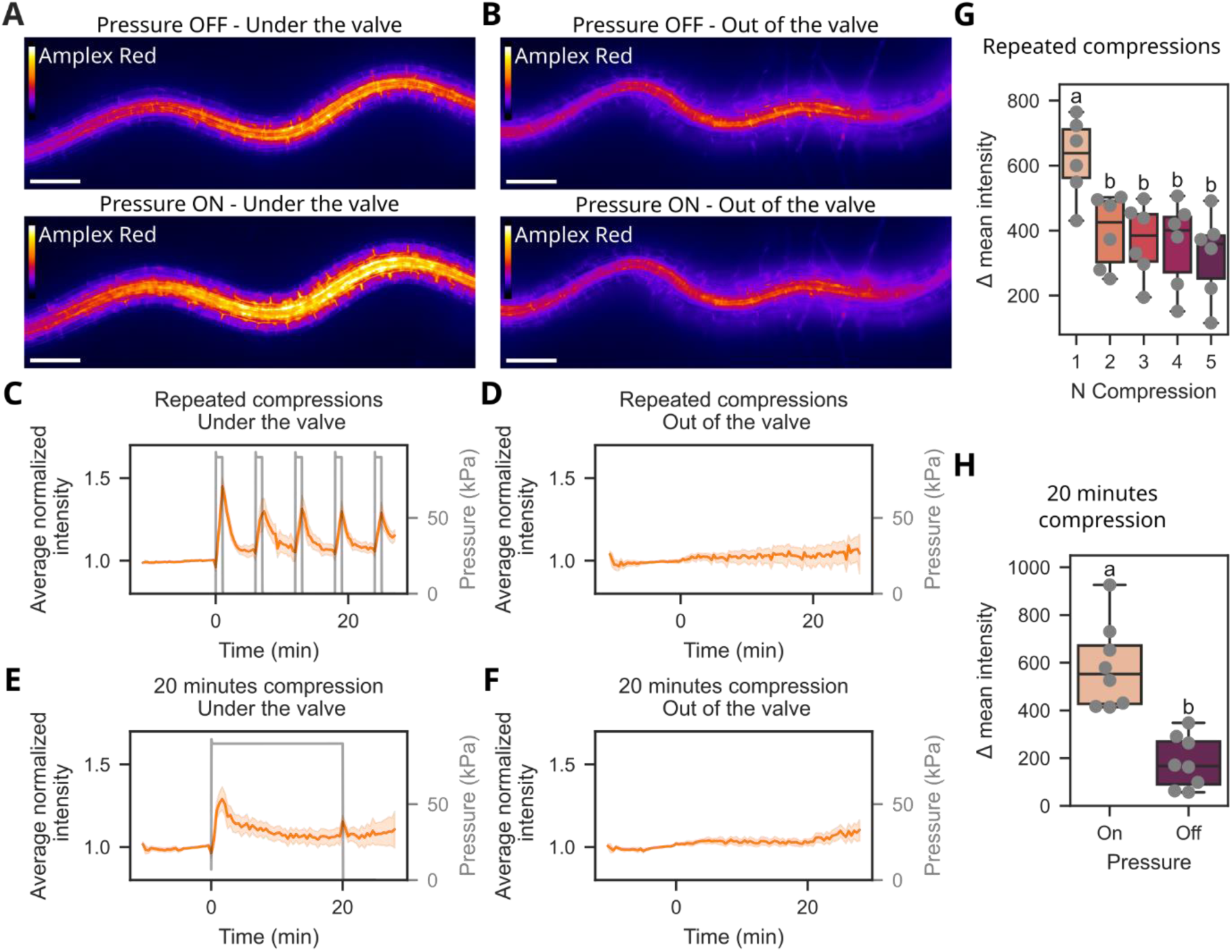
The H_2_O_2_ response to compression is transient and local. H_2_O_2_ probe intensity under the valve before and during compression with 90 kPa pressurized air injected in the pressure channel **(A)** or out of the valve for the same root and times **(B)**. Same Amplex Red intensity color scale for both images. 100 µm scale bar. **(C)-(F)** Mean of averaged normalized Amplex Red intensity of roots under the valve **(C), (E)** or out of the valve **(D), (F)** for 5 repeats of a 1-minute short compression **(C) and (D)** or 20 minutes long compression for other roots **(E) and (F)**. Time origin corresponds to the beginning of the pressure protocol. Pressure values applied in the pressure channels are shown in gray. **(G)** Amplitudes of the H_2_O_2_ response for each compression repeat. **(C,D,G)** N = 3 experiments, n = 6 positions for 3 different roots. Different letters indicate significant differences: p-value < 0.05, repeated measures ANOVA and post-hoc paired T-tests. **(H)** Amplitudes of H_2_O_2_ response at the onset (Pressure On) and at the release of pressure (Pressure Off). **(E,F,H)** N = 5 experiments, n = 8 positions for 5 different roots. Different letters indicate significant difference, p-value < 0.05, paired T-test.

Upon repeated compressions, the amplitude of the Amplex red signal decreased significantly by 36 % between the first and second compression and remained at a low level upon further stimulation (Fig. 2C and G, Supplementary Figure 4). This indicates an attenuation of the H_2_O_2_ response to root compression. This decrease could not be due to Amplex red depletion, as it was continuously replenished by the flow system.

To test whether the H_2_O_2_ accumulation was reversible even when compression was maintained, we applied a long compression of 20 minutes. This triggered local transient increases of Amplex Red signal both at the onset and at the release of pressure (Fig. 2E and F, Supplementary Figure 4). H_2_O_2_ accumulation reached a maximum 89 ± 24 s after pressure started, and the mean fluorescence intensity returned to its initial value within approximately 15 minutes. Releasing the pressure also induced a transient H_2_O_2_ accumulation reaching a maximum after 3.0 ± 3.4 s. The signal observed at the pressure release had a significantly lower amplitude compared to the one at the onset of pressure (Fig. 2H). This suggests that H_2_O_2_ accumulates in response to root deformation rather than to root compression *per se*, as root deformation happens both at the pressure onset and release. Taken together, the results obtained when measuring H_2_O_2_ accumulation with Amplex red show signal attenuation, responses at both pressure onset and release and timescales of the response similar to what was observed for the calcium response in previous work. This raises the question of the relationships between the calcium and H_2_O_2_ responses.

### H_2_O_2_ and calcium responses occur in three successive kinetic phases

To better understand the relationships between the calcium and H_2_O_2_ signals, we first examined their temporal relationships by monitoring the two responses on the same root. For this, we used roots expressing GCAMP calcium probe continuously perfused with with 10 µM Amplex Red. Sequential excitation acquisition was performed in wide-field microscopy on the root zone under the rootchip valve. The response to a 1-minute compression was monitored with a time resolution of 1 frame every 2 to 6 seconds (Fig. 3A).

**Figure 3:**
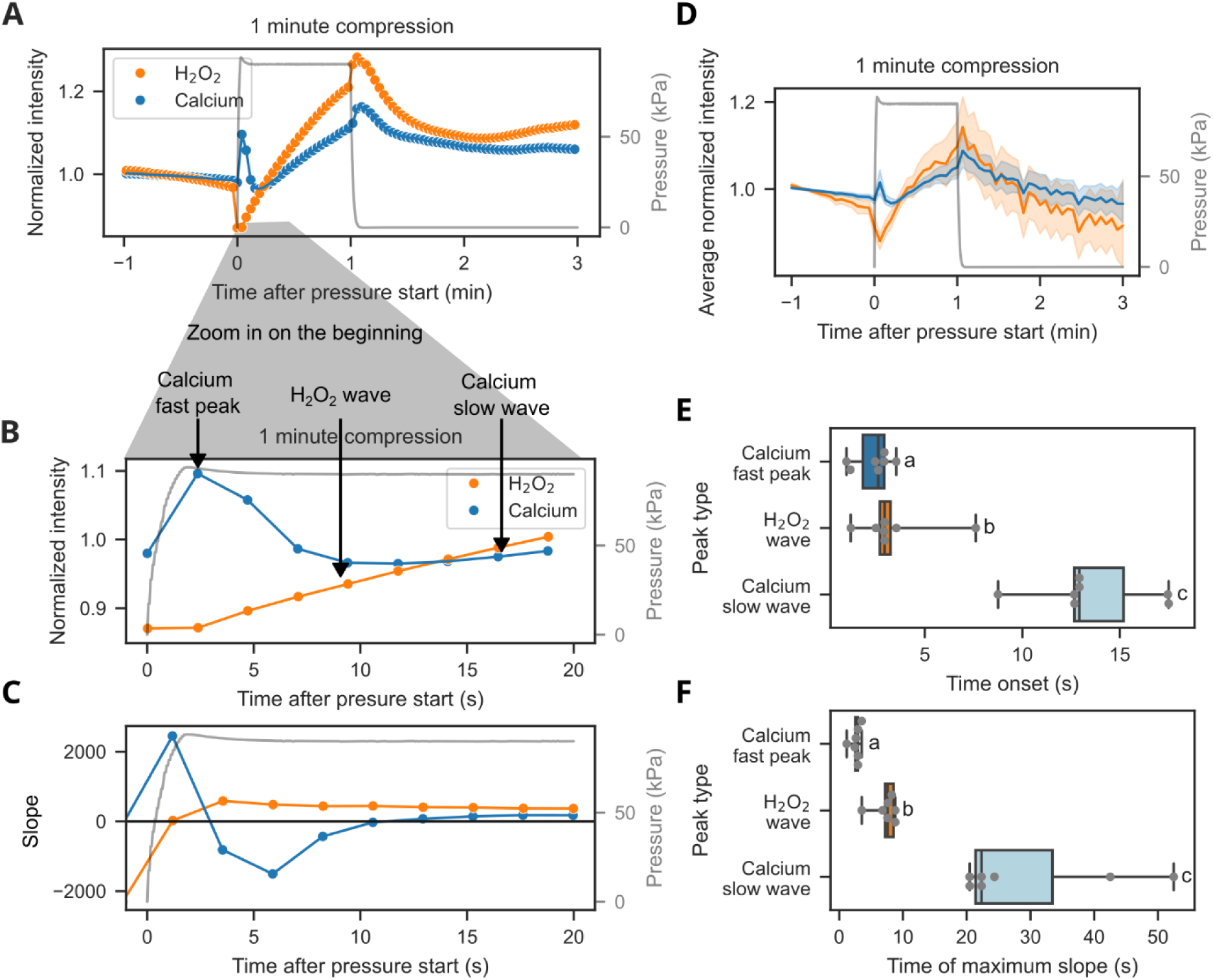
Kinetic phases of H_2_O_2_ and calcium responses to a compression. **(A)** Normalized fluorescence intensity of H_2_O_2_ and calcium signals (Amplex Red in orange and GCAMP in blue respectively) during a 1-minute compression on one root. **(B)** Zoom in on the first seconds after the compression started. Three kinetic phases can be identified (black arrows): the calcium fast peak, the H_2_O_2_ wave and the calcium slow wave. **(C)** Slope of the mean intensity curve calculated for each pair of consecutive timepoints. The beginning of a peak or wave is defined as the time point when the slope becomes positive. **(D)** Average of the normalized intensity and confidence interval at 95%. N = 4 experiments, n = 7 positions on 7 different roots. **(E)** Time of beginning for each kinetic phase in different roots. **(F)** Time at which the maximum slope value is reached.Different letters indicate significant difference, p-value < 0.05, Friedman test and pairwise Wilcoxon tests. N = 4 experiments, n = 7 positions on 7 different roots.

A first overview of mean fluorescence intensities over time revealed a biphasic response for cytosolic calcium and a monophasic response for H_2_O_2_ (Fig. 3A, B, D, Supplementary Figure 5). Hereafter, these phases will be referred to as the calcium fast peak for the first phase of calcium response, the calcium slow wave for the second phase and the H_2_O_2_ wave for the H_2_O_2_ response, as shown in Figure 3B. To better characterize these kinetic phases, the slope of the mean intensity variation was calculated as a function of time (shown in Fig. 3C for one root). This allowed determining for each root: the onset time of a phase, as the time point when the slope becomes positive; the slope value at the onset; the maximum slope and the time point when this maximum is reached for each phase (Fig. 3C, E, F Supplemental Fig. 6 A, B).

For the calcium fast peak, an increase happened within 4 seconds after the onset of pressure application (Fig. 3 E), followed by a fast decrease. The fast calcium peak rose significantly earlier (Fig. 3 E) and faster than the calcium slow wave (Fig. 3 E,F). The slow calcium wave started between 8.8 and 17.5 seconds and reached maximum slope values about 5 times lower than the first calcium peak (Supplemental Fig. 6), suggesting different mechanisms of calcium signaling.

The H_2_O_2_ wave started within 8 seconds (3.4 ± 2 s) after the onset of pressure, at the same time or just after the beginning of the calcium fast peak and significantly earlier than the calcium slow wave (Fig. 3E, F, Supplementary Fig. 5). The H_2_O_2_ wave shares long-lasting features with the calcium slow wave but does not exactly follow the same kinetics, with an earlier start and faster kinetics.

To summarize, we could characterize 3 kinetic phases of response to lateral compression. (1) A calcium fast peak starting a few seconds after pressure application, with the fastest increase and lasting only for a few seconds. (2) A long-lasting H_2_O_2_ wave starting almost at the same time as the calcium fast peak, with a slow increase at the beginning that accelerates to reach a maximum speed about 3 seconds later. (3) A long-lasting calcium slow wave, starting after the H_2_O_2_ wave and with a much slower increase compared to the two other events. This suggests a scenario in which the fast calcium peak and the H_2_O_2_ wave precede the calcium slow wave. The calcium slow wave could therefore depend on the H_2_O_2_ wave and/or the calcium fast peak.

### The H_2_O_2_ response is ubiquitous in the root while Ca responses localize in the root center

To further characterize the interplay between H_2_O_2_ and calcium responses to root compression, we analyzed their spatial relationships. To resolve the different signals at the tissue level, we performed confocal cross-section imaging using roots expressing GCAMP and stained with Amplex Red. In these cross sections (Fig. 4A, Fig. 1C), the fluorescence intensity was high in the peripheral root tissues, but the signal attenuated in the central root tissues, due to light absorption by the tissues. We defined 3 regions of interest: the epidermis, the cortex and the root center (Fig. 1C).

**Figure 4:**
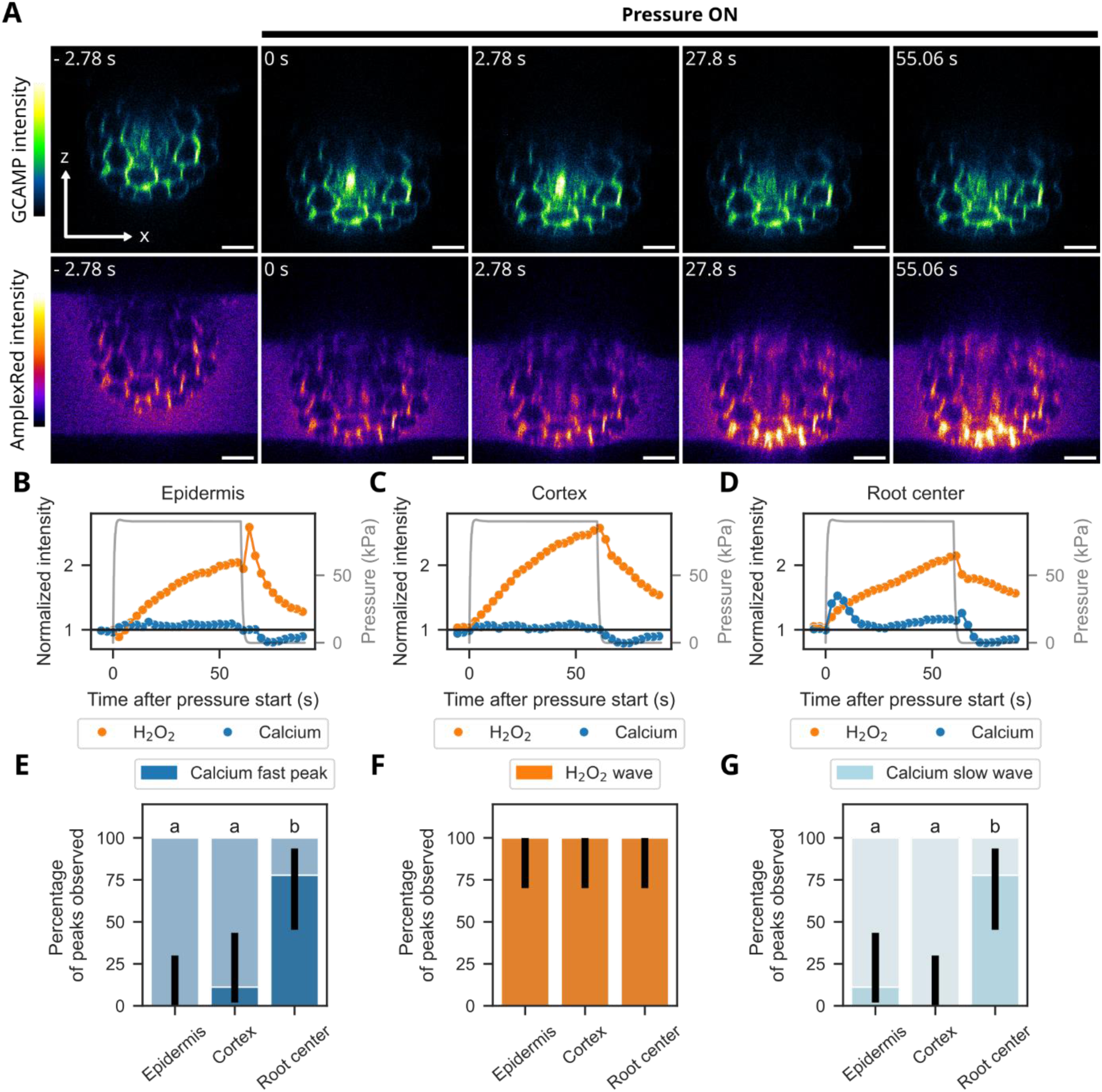
Localization of H_2_O_2_ and Calcium responses in the root tissues. **(A)** Confocal cross sections of a root before and during a 1-minute compression, with color coded fluorescence intensity of the GCAMP calcium probe (top) and Amplex Red H_2_O_2_ probe (bottom). Frames shown here are extracted from a timelapse acquisition, timestep 2.78s. Time origin at the onset of the pressure application. **(B)**, **(C)** and **(D)** Ratio between the mean fluorescence intensity and the baseline intensity of H_2_O_2_ and calcium probes, in orange and blue respectively. Baseline intensity is the mean of fluorescence intensity before pressure. This ratio is calculated on ROIs corresponding to the epidermis **(B)**, the cortex **(C)** and the root center **(D)**. The curves shown in B, C and D are the responses in the root shown in **(A)**. **(E)**, **(F)**, **(G)** Percentage of roots that responded exhibiting **(E)** a calcium fast peak, **(F),** a H_2_O_2_ wave and **(G)** a calcium slow wave observed in the epidermis, cortex or root center for N = 3 experiments and n = 9 positions on 9 different roots. Confidence intervals of the percentages are shown with black lines (Wilson method). Different letters indicate significant differences, p-value < 0.05, χ² independence test and pairwise χ² with Benjamini-Hochberg correction.

Generally, the responses to a 1-minute compression observed with confocal cross-section acquisitions followed the same kinetics as described previously with wide-field acquisition (Supplementary video 2). Figure 4A shows an initial sharp increase in GCAMP signal less than 3 s after pressure onset. This calcium fast peak was followed by Amplex Red signal increase visible after 25 s and another GCAMP slower increase visible towards the end of pressure application, after 55 s. The Amplex Red signal was not homogeneous in the root, with higher fluorescence values in the most external tissues, probably due to its integration in the root from the external medium. GCAMP also presented different basal values of fluorescence intensity depending on the tissue. To compare the signal in the different zones, the mean fluorescence intensity was normalized by the intensity value before pressure (referred to as baseline) for each region of interest (Fig. 1C; Fig. 4 B, C, D).

The H_2_O_2_ wave was observed in the epidermis (Fig. 4B), the cortex (Fig. 4C) and the root center (Fig. 4D). No significant differences were detected between the percentages of H_2_O_2_ waves observed in these regions among all tested roots (Fig. 4F). For the root shown in Figure 4A and quantified in Figure 4 B-D, the calcium fast peak and calcium slow wave were visible only in the root center. The percentage of calcium fast peaks or calcium slow waves observed among all tested roots confirms a much higher incidence of the calcium response in the root center compared to the epidermis and cortex (Fig. 4E, G, Supplementary Fig. 7 and 8). Similar results for the calcium response were obtained with the ratiometric RGECO-mTurquoise probe, confirming that the differences in calcium response are not due to differences in GCAMP expression levels between tissues (Supplementary Fig. 9). This indicates that the H_2_O_2_ response to a compression is ubiquitous in the root tissues, whereas the calcium biphasic response is mainly localized in the root center.

The kinetics experiments showed successive phases that could be triggered by one another, but localization experiments revealed that H_2_O_2_ increases in tissues where no calcium response is observed. We thus wonder whether a causal relationship exists between the H_2_O_2_ and the calcium responses.

### Causal links between H_2_O_2_ and Calcium in response to root compression

To decipher the potential causal links between H_2_O_2_ and calcium responses, we performed inhibition experiments. To limit external calcium influx through calcium-permeable channels, we omitted calcium in the medium and added 1 mM Gd^3+^, a broad spectrum inhibitor of calcium channels, a condition hereafter referred to as 0 Ca^2+^ + Gd^3+^ (Ermakov et al., 2010; Tran et al., 2017). The production of ROS by NADPH oxidases was inhibited by DPI at 50 µM or 100 µM (Morré, 2002; Foreman et al., 2003; Reis et al., 2020). GCAMP and Amplex Red signals were imaged in treated roots in wide-field microscopy during short compressions of 1 or 5 minutes, and the effects on the 3 kinetic phases observed previously were assessed.

Upon external calcium removal and mechanosensitive channels inhibition, the calcium fast peak disappeared from the GCAMP fluorescence intensity curves (blue curve for normalized intensity average for different roots and round markers for one representative root in Fig. 5A). Overall, the percentage of calcium fast peak observed in 0 Ca^2+^ + Gd^3+^ roots was 10 times lower than in the control or DPI treated roots (Fig. 5G). In contrast, ROS production inhibition had no significant effect on the fast calcium peak. These results indicate that the calcium fast peak requires influx of external calcium, potentially via mechanosensitive channels.

**Figure 5:**
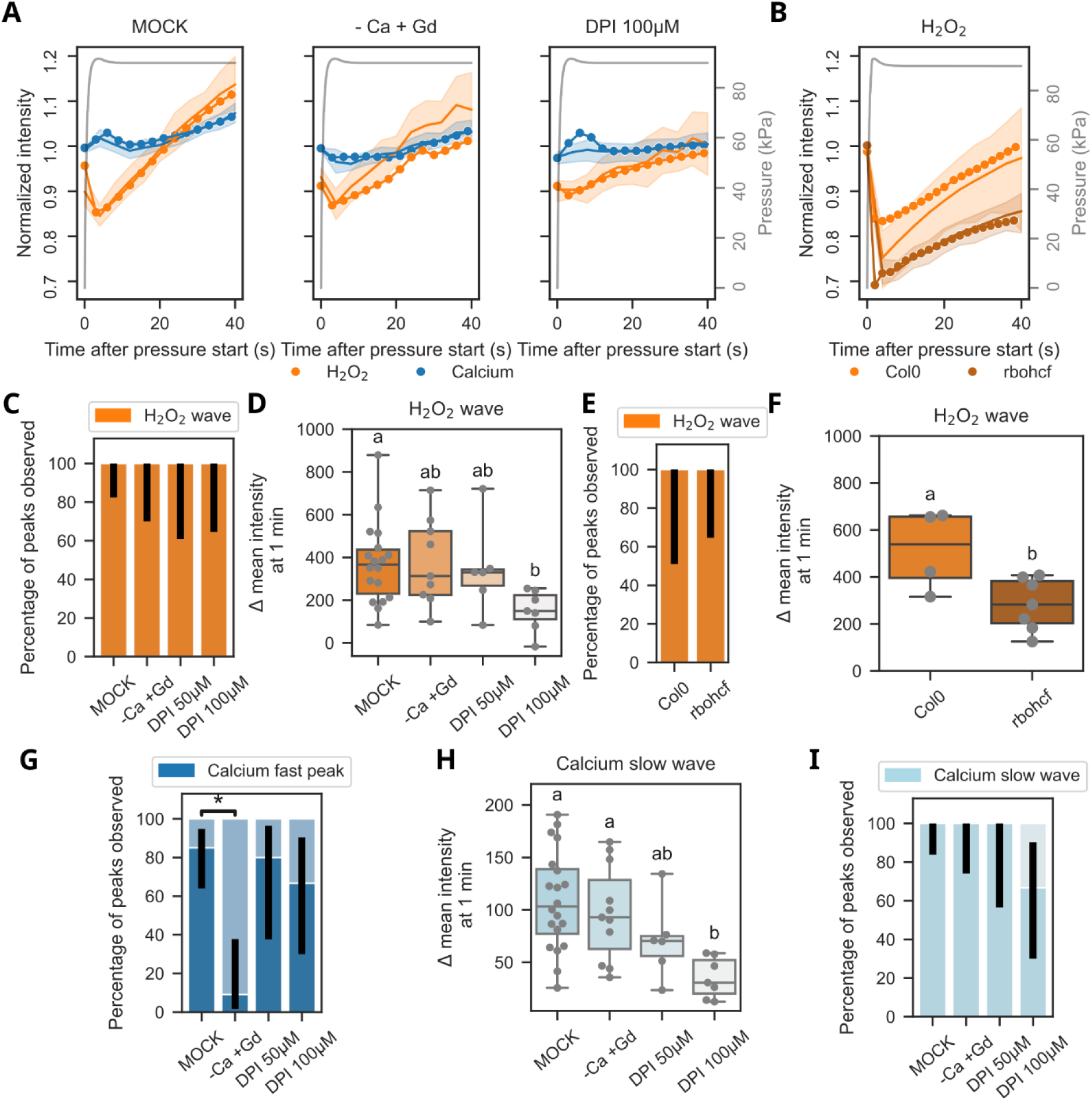
Effects of ROS production or mechanosensitive channels inhibition on H_2_O_2_ and calcium response phases. **(A)** Representative examples of normalized intensity (round markers) and average normalized intensity (line and filled zone, 95% confidence interval) of Amplex Red H_2_O_2_ probe in orange and GCAMP calcium probe in blue, during a 1-minute compression of roots incubated in Hoagland medium (MOCK), Hoagland 0 calcium medium with 1 mM Gd^2+^ (-Ca +Gd) or Hoagland medium with 100 µM DPI (100µM DPI). The calcium fast peak is not visible for -Ca +Gd condition. (**B)** Representative examples of normalized intensity (round markers) and average normalized intensity (line and 95% confidence interval) of Amplex Red H_2_O_2_ probe in WT and rbohcf roots, in orange and black respectively. **(C,D)** Percentage of H_2_O_2_ waves observed for control roots (MOCK) or roots treated with 0 mM calcium and 1mM Gd^2+^, 50 µM DPI or 100 µM DPI **(C)** or for WT and rbohcf mutant roots **(D)**. Amplitude at 1 minute of the H_2_O_2_ wave **(E)** or the calcium slow wave **(H)**,for control roots or roots treated with 0 mM calcium and 1 mM Gd^2+^, 50 µM DPI or 100 µM DPI. Different letters indicate significant differences, p-value < 0.05, Welch ANOVA and pairwise Games Howell tests**. (F)** Amplitude at 1 minute of the H_2_O_2_ wave for WT and rbohcf roots. Different letters indicate significant differences, p-value < 0.05, Mann-Whitney U test **(G) (I)** Percentage of calcium fast peaks (G) or calcium slow wave (I) observed for control roots (MOCK) or roots treated with 0 mM calcium and 1mM Gd^2+^, 50 µM DPI or 100 µM DPI. **(C) (D) (G) (I)** Confidence intervals are shown in black (Wilson method). Star indicates significant difference, p-value < 0.05, χ² independence test and pairwise χ² with Benjamini-Hochberg correction. (C) (G) (I) and (E) (H) N ≥ 3, n ≥ 5 positions on n ≥ 5 different roots for each treatment. (D) and (F) N ≥ 2, n ≥ 4 positions on 4 roots.

In contrast, external calcium removal and mechanosensitive channel inhibition had no significant effect on the features of the H_2_O_2_ wave or the calcium slow wave (Fig. 5C,D and H,I). This suggests that the calcium fast peak, inhibited by the 0 Ca^2+^ + Gd^3+^, is not required for the H_2_O_2_ wave and calcium slow wave.

The H_2_O_2_ wave was affected by inhibition of ROS production with 100 µM DPI. This treatment reduced the amplitude of the H_2_O_2_ wave by 59 % compared to control roots (Fig. 5 D), while it had no effect on the percentage of H_2_O_2_ waves observed (Fig. 5C). NADPH oxidase inhibition did not induce a complete loss of the ROS response but a slower response, reaching lower intensity values. This indicates that NADPH oxidases activity contributes to but does not fully account for ROS production in response to root compression. Similar results were observed for the calcium slow wave, with significantly reduced amplitudes and maximum slopes after 100 µM DPI treatment but no significant effect on the percentage of peaks observed (Fig. 5 H,I). This suggests that the calcium slow wave partially depends on ROS production by NADPH oxidase. To confirm that H_2_O_2_ accumulation relies on NADPH oxidase activity, we monitored the H_2_O_2_ response in the *rbohc/f* double mutant lacking the two major isoforms of this enzyme expressed in roots. We observed that the amplitude of the H_2_O_2_ response was decreased by about 40% in the double mutant, while the frequency of occurrence of the response was not significantly affected (Fig. 5B, E, F, supplementary Fig. 10).

### Molecular players underlying calcium and H_2_O_2_ responses

The mechanosensor involved in the fast calcium response is most likely a mechanosensitive channel residing on the plasma membrane. The lack of causal link between the fast calcium peak and the subsequent H_2_O_2_ response and slow calcium response suggests that the slow responses may rely on another mechanoperception mechanism. To test whether the receptor kinase FERONIA, a potential mechanoreceptor mediating many responses to mechanical stimuli is involved, we monitored the calcium and H_2_O_2_ responses in the *fer4* mutant expressing GCAMP (Duan et al., 2010; Shih et al., 2014; Malivert and Hamant, 2023). The mutation did not affect the frequency of occurrence of fast calcium peaks (fig. 6C). In contrast, we observed strong decreases in the amplitude of both H_2_O_2_ and slow calcium responses in *fer4* (Fig. 6A, B, F, G), while the frequency of occurrence of the responses was not significantly affected (Fig. 6D, E).

**Figure 6:**
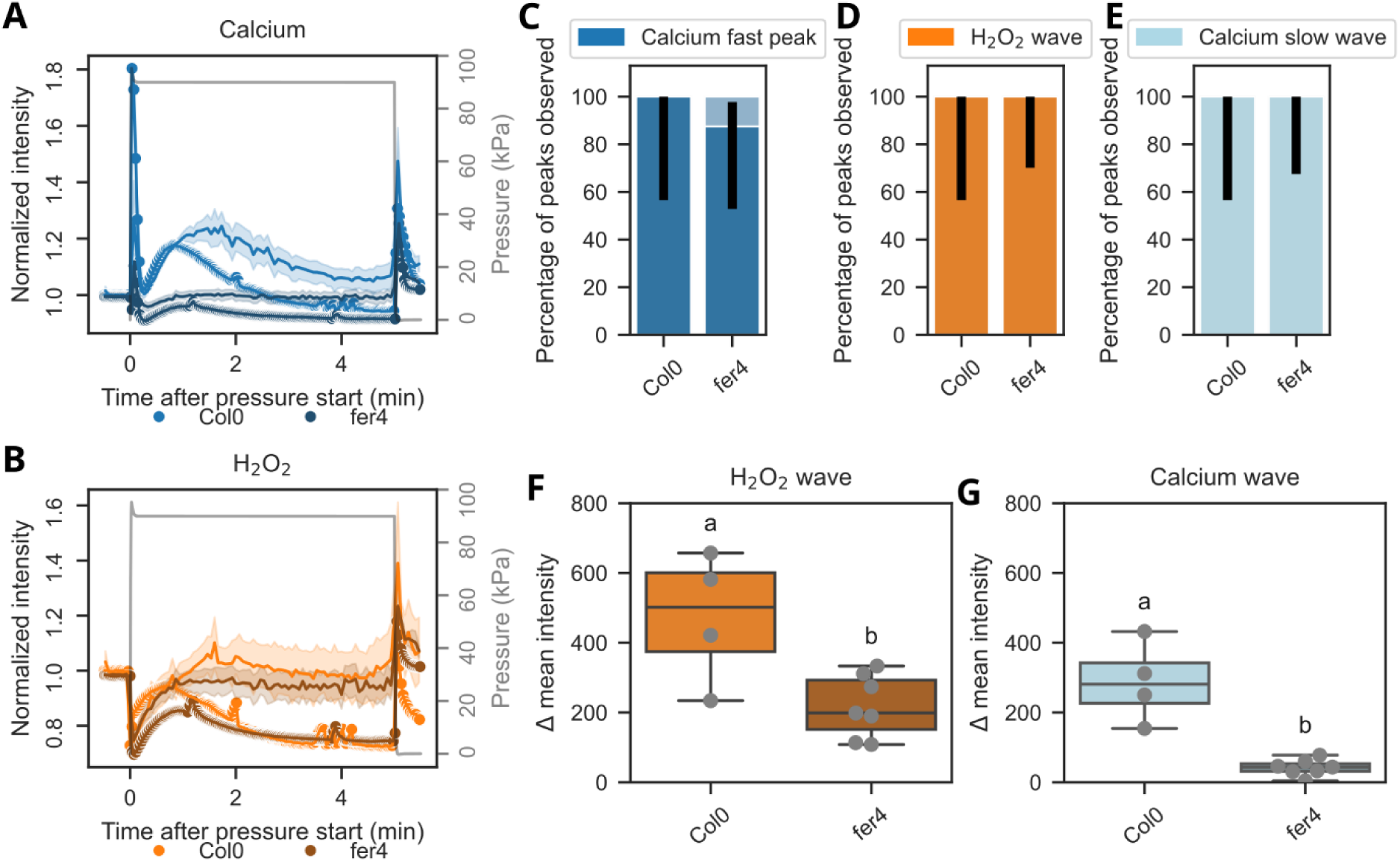
Effect of FERONIA loss of function on H_2_O_2_ and calcium responses. **(A) (B)** Representative examples of normalized intensity (round markers) and average normalized intensity of Amplex Red H_2_O_2_ probe in orange and GCAMP calcium probe in blue, during a 1 minute compression of WT **(A)** or fer4 mutants **(B)**. Percentage of calcium fast peaks **(C)**, H_2_O_2_ waves **(D)** and calcium slow waves **(E)** observed in WT or fer4 mutant roots. Confidence intervals are shown in black (Wilson method). N.S. χ² independence test and pairwise χ² with Benjamini-Hochberg correction. N ≥ 2, n ≥ 5 positions on 5 different roots. Amplitude of the H_2_O_2_ wave **(F)** and the calcium slow wave **(G)** in WT and fer4 mutant roots. Different letters indicate significant differences, p-value < 0.05, Mann-Whitney U test. N ≥ 2, n ≥ 5 positions on 5 different roots.

To conclude, two mechanisms could be involved in the H_2_O_2_ and calcium responses to compression: (1) a mechanism relying on external calcium and mechanosensitive channels triggering a calcium fast peak and (2) a H_2_O_2_ wave, partially dependent on ROS production by NADPH oxidase and FERONIA mechanoperception, linked to a slower calcium wave. As the slow calcium wave is not affected by the lack of external calcium, this phase may rely on calcium release in the cytosol from intracellular sources such as the endoplasmic reticulum or the vacuole.

## Discussion

### Two mechanisms of mechanotransduction in response to root compression

In this work, we show that root lateral compression induces H_2_O_2_ transient accumulation within the root and analyze its relationships with the previously reported biphasic calcium response (Audemar *et al*., 2023). The three events occur sequentially, starting with the fast calcium peak, then the H_2_O_2_ response and finally the slow calcium wave. Interestingly, although the epidermal and cortical cells undergo the highest deformation, the calcium response occurs in the root center. Based on kinetic, pharmacological and genetic evidence, we propose a model for the interplay between H_2_O_2_ and calcium responses and the transduction of a root lateral compression into these primary responses (Fig. 6). We highlight two distinct mechanisms. The first one, responsible for the cytosolic calcium fast peak, is dependent on external calcium entry through mechanosensitive channels. The second one, comprising the H_2_O_2_ wave and the cytosolic calcium slow wave partially relies on mechanoperception by FERONIA and the activity of NADPH oxidases such as RBOHC or F.

**Figure 7:**
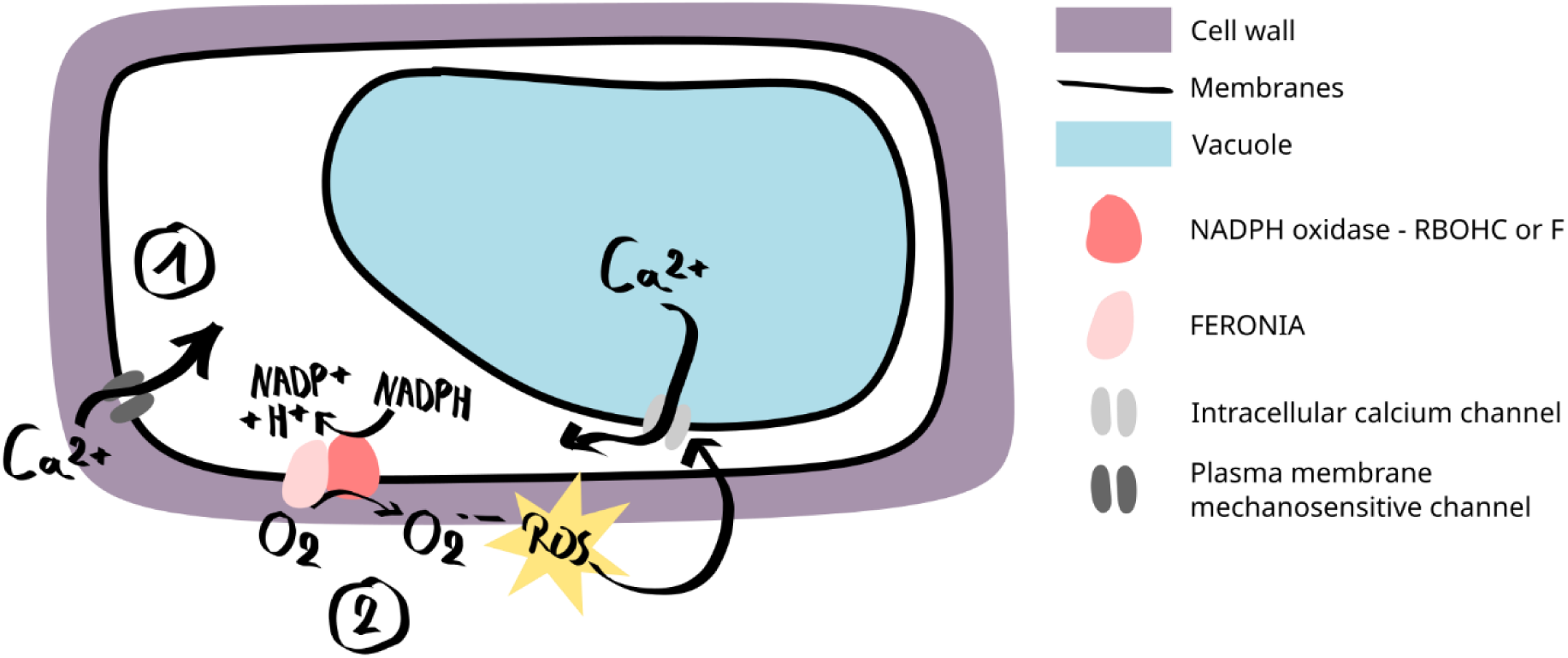
Two mechanisms of mechanotransduction at the origin of Calcium and H_2_O_2_ responses to lateral compression. The first mechanism (1) corresponds to external calcium entry via mechanosensitive channels, triggering the fast calcium peak. The second mechanism (2) involves FERONIA, potentially activating the NADPH oxidases RBOHC and/or RBOHF. This would induce ROS accumulation followed by ROS-dependent calcium release from intracellular sources, possibly from the vacuole.

### Attenuation of H_2_O_2_ and Ca^2+^ responses

We observed an attenuation of the H_2_O_2_ response upon repeated compressions, as observed for the calcium slow wave by Audemar *et al*. The attenuation could be an intrinsic property of the mechanotransduction machinery involved in the ROS accumulation. Attenuation has been observed in various mechanotransduction responses. For example, early mechanoresponsive genes show a lower increase in expression after a second bending of the poplar stem (Martin et al., 2010; Pomiès et al., 2017). In *Mimosa pudica*, a first touch induces full leaf folding, but the frequency of partial leaf folding increases with touch repetitions (Amador-Vargas et al., 2014). In *Arabidopsis* protoplasts, the activity of the calcium-permeable force-gated channel Rapid Mechanically Activated (RMA) decreases upon repeated stimulations (Guerringue et al., 2026). Attenuation of the responses to repeated mechanical stimuli is thus observed at different scales and is a broad phenomenon that could help plant adaptation to recurrent stresses in their environment.

### Plasma membrane MS channels underlying the fast calcium peak

The calcium fast peak has similar kinetics as the initial calcium response to touch observed in pavement cells and the first phase of calcium response to root bending (Shih et al., 2014; Howell et al., 2023). This points toward fast and transient calcium increase as a common feature of plant responses to mechanical stimulation. The fast entry of external calcium, inducing the calcium fast peak, could be due to plasma membrane mechanosensitive (MS) channel opening/activation. The fast activation and deactivation kinetics of mechanosensitive channels fit well with this hypothesis. Activation kinetics of MS channels are in the few milliseconds range and deactivation in a few seconds range (Frachisse et al., 2020). Among MS channels, the calcium permeable conductance, RMA is a good candidate to mediate the fast calcium response but further molecular characterization is needed to uncover its potential role (Guerringue et al., 2026). Other candidates are the stretch-activated Ca^2+^ permeable MCA1 and MCA2 channels or the OSCA channels family, involved in calcium signaling in response to osmotic stress (Yoshimura et al., 2021; Pei et al., 2024). The MSL anionic channels could also play an indirect role, especially MSL10 which is involved in the rapid and transient cytosolic calcium increase observed in response to cell swelling (Basu and Haswell, 2020). Other non-mechanosensitive calcium channels sensitive to Gd^3+^ could be involved, such as the CNGCs (Tan et al., 2020). Further experiments with mutants for these channel candidates need to be conducted to conclude on the molecular players inducing the calcium fast peak.

### Mechanically triggered ROS production induces calcium release from intracellular sources

Our results indicate that inhibiting the fast calcium response does not influence the slower rising and long-lasting H_2_O_2_ and calcium responses. These responses partially require NADPH oxidase activity and are not impacted by external calcium depletion. We showed that a second mechanotransduction mechanism, involving FERONIA, triggers ROS production by NADPH oxidases RBOHC and F and a ROS-induced calcium release from intracellular sources. The involvement of NADPH oxidase and the reversibility of the signal by fast washing-out of the resorufin both point to mostly extracellular H_2_O_2_ production. The fraction of the H_2_O_2_ response that is not sensitive to DPI could be linked to intracellular production. The finding that an H_2_O_2_ response is still visible in the *rbohc/f* mutant may indicate that other NADPH oxidase isoforms are involved.

Several reports demonstrated the importance of mechanosensitive ion channel activation upstream of ROS accumulation upon mechanical stimulation. Recent results showed a role of MCA1 MS channel in apoplastic H_2_O_2_ accumulation in response to root bending (Kulich et al., 2026). In addition, ROS accumulation induced by cell swelling required MSL10 activity (Basu and Haswell, 2020). NADPH oxidase activity can be regulated by calcium binding and calcium dependent protein kinases and these processes are thought to be involved in ROS and calcium waves propagation in response to salt stress or hypoxia-induced ROS production for example (Ogasawara et al., 2008; Evans et al., 2016; Yu et al., 2024).

However, our experiments using Gd^3+^ indicate that the mechanotransduction mechanism inducing NADPH oxidase activation in response to root compression is independent of the preceding calcium fast peak. Our results with *fer4* mutants suggest that FERONIA is involved in this independent mechanotransduction mechanism. FERONIA (FER) is a cell-wall binding receptor-like kinase (RLK) from the *Catharanthus roseus* receptor-like kinase 1 like (*Cr*RLK1L) family involved in cell wall integrity sensing and various cellular processes (Malivert and Hamant, 2023). FER has been shown to regulate NADPH oxidase-dependent ROS accumulation in roots (Duan et al., 2010; Lu et al., 2024). Smokvarska *et al*. also showed that FER is necessary to induce ROS accumulation in roots under hyperosmotic conditions (Smokvarska et al., 2023). Upon root bending, Shih *et al*. reported a biphasic calcium response on the convex side of the root in the same time scale as the calcium fast peak and calcium slow wave observed in our root compression experiments (Shih et al., 2014). In *fer* loss-of-function mutants, the second calcium phase was lost, similar to the effect on the calcium slow response we observed in our experiments. Further experiments should be done to confirm whether FERONIA induces RBOH C and F activation in our conditions.

We observed that the second calcium wave initiated a few seconds after the H_2_O_2_ response and was dependent on ROS production. This suggests the involvement of ROS-activated calcium permeable channels. Such channels were first identified at the plasma membrane in guard cells and in root epidermal cells (Pei et al., 2000; Demidchik et al., 2003). However, we found that the calcium slow wave is not affected by the depletion of external calcium and inhibition of plasma membrane channels. This suggests that it involves the release of calcium from intracellular sources such as the endoplasmic reticulum or the vacuole. Annexin 1 is a phospholipid-binding protein that could form atypical membrane channels in the ER and induce sustained calcium elevation in response to ROS (Richards et al., 2014; Tichá et al., 2020). Further experiments would be needed to test the role of Annexin1 or identify other channels regulated by ROS in response to root compression.

We observed a partial inhibition of the H_2_O_2_ response when inhibiting NADPH oxidases and in *rbohc/f* mutants. This suggests that either 100 µM DPI does not inhibit NADPH oxidases completely or that another source of H_2_O_2_ accumulation is involved in addition to NADPH oxidases. In the *rbohc/f* mutant, there could be a redundancy between other RBOH family members. H_2_O_2_ accumulation can be linked to an increase in production and/or a decrease in antioxidant activity. As the ROS wave was not fully inhibited in our experiments, it is not possible to know if the calcium slow wave is fully dependent on ROS.

### Localization of the responses

In our previous characterization of the root compression induced in the microfluidic valve rootchip, we showed that the root center was less deformed than the outer layers, especially the epidermis (Audemar et al., 2023). Here, we find that upon long compression ROS and calcium responses occur at the onset and release of pressure. This suggests that ROS and calcium respond to the organ/cell deformation. The calcium increases were observed mainly in the root center, the less deformed zone, while the ROS response was observed in the whole root. This localization could be explained by differences in mechanoreceptors and calcium channels expression patterns or by different mechanical properties of the tissues, facilitating or hindering mechanotransduction.

In Arabidopsis primary root, MCA1 and MCA2 are preferentially expressed in the vascular tissues (Yamanaka et al., 2010). Information about other MS or calcium channels localization in root tissues is missing, however members of the main MS channels families seems mainly expressed in the center of the root (Hartmann et al., 2021). The broad localization of the ROS response is consistent with the broad expression pattern of Feronia in Arabidopsis roots (Dong et al., 2019).

The cells from the central cylinder are expected to be more rigid because of wall lignification. However, the mechanotransduction initiates in this region, suggesting that only a minimal deformation is sufficient to trigger a response. Information on the mechanical properties of the different root tissue layers is scarce and the relationship between membrane tension and cell wall mechanical properties remains poorly understood (Alonso Baez et al., 2026). Fino *et al*. showed that pericycle cells have a specific mechanical environment, as pericycle-endodermis interface walls are softer than walls of the outer layer interfaces and different membrane properties are observed (Fino et al., 2025). It would be interesting to investigate the calcium response in the root center with higher imaging resolution to test whether pericycle cells are involved in the calcium response to root compression.

## Materials and methods

### Plant material and growth conditions

All the lines used are in the *Arabidopsis thaliana* Col0 background. *GCAMP* (*pUB10::GCAMP3*) calcium probe seeds from (Krogman et al., 2020) and *fer4 GCAMP* were kindly provided by Herman Höfte and Stéphanie Afonso. *RGECO-mTurquoise* (*pUB10::RGECO1-mTurquoise*) ratiometric calcium probe seeds are described in (Waadt et al., 2017). *rbohc/f* mutant seeds were kindly provided by Ivan Kulich and Jiri Fiml and are described in (Kulich et al., 2026). Seeds were sterilized for 5 minutes in ethanol 70% + 0.05% SDS (Sodium Dodecyl Sulfate) and rinsed twice for 5 minutes in ethanol 96%. To prepare seedlings for the insertion in the rootchip, pipette tips were filled with Hoagland medium with 1% phyto-agar (1.5 mM Ca(NO_2_)_2_, 0.28 mM KH_2_PO_4_, 0.75 mM MgSO_4_, 1.25 mM KNO_3_, 0.5 µM CuSO_4_, 1 µM ZnSO_4_, 5 µM MnSO_4_, 25 µM H_3_BO_3_, 0.1 µM Na_2_MoO_4_, 50 µM KCl, 3 mM MES, 10 µM Fe-HBED, pH 5.7). Cones were then cut, and the tips were placed in a petri dish filled with the same medium. Seeds were sown on each cone with a toothpick. After 3 days of stratification (4°C dark), seeds were incubated for 4 days at 23°C (16 h light/8 h dark). Fluorescent seedlings were then selected with a macroscope (AZ 100, Nikon) and transferred in the microfluidic valve rootchip. The rootchips were manufactured as described previously in (Audemar et al., 2023). In the rootchip, seedlings grew for 3 days in the same temperature and light conditions with a tilting of 45°, allowing the primary root to enter the root channels. These channels were connected to syringes and syringe pumps, refreshing the liquid Hoagland medium with a 1 µL/minute flow rate (WPI, AL-1000).

For H_2_O_2_ observation, roots were incubated with Hoagland medium containing 10 µM Amplex Red (A12222, ThermoFischer) for 1 h before the acquisition and during the acquisition, at a flow rate of 4 µL/minute. To inhibit ROS production, DPI (Diphenyleneiodonium, CAS 4673-26-1, Merck, stock solution 2mM in water) was added at 50 or 100 µM at the incubation step. Hoagland medium without calcium (0 mM Ca(NO_2_)_2_) and with 1 mM Gd^2+^ (Gadolinium nitrate hexahydrate, stock solution 1M in water) was used for mechanosensitive channels inhibition experiments.

### Image acquisition

After a total of 7 days of incubation, roots had grown past the valve (i.e. the deformable PDMS layer between pressure and root channels) and could be observed under the microscope. Rootchips were connected to new syringes filled with media containing Amplex Red, Amplex Red + DPI or Hoagland 0 calcium, Amplex Red + Gd^2+^ and placed in a 3D printed sample holder located in a standard 96-well plate holder. Syringe pumps imposed a flow rate of 4 µL/minute in the root channels during the incubation and the experiment. To inject pressurized air into the pressured channels and deform the valve above the root, pressure channels were connected to a pressure box (MFCS-EZ, Fluigent). Pressure protocols were designed and applied via the MAT software (Microfluidics Automation Tool, Fluigent). For short compressions, the protocol was 1 minute or 5 minutes of pressure at 90 kPa followed by 5 minutes at 0 kPa. This sequence was repeated 5 times in the repeated compressions protocol. For long compressions, pressure at 90 kPa was applied for 20 minutes.

To image the global response on the root zone under the valve, a Leica DMI100 wide-field inverted microscope was used, with a quad band dichroic block and emission filter (dichroic block BP 380-410, 475-500, 545-565, 625-650 nm Chroma; emission filter BP 425-465, 510-530, 580-620, 670-740 nm). When imaging intracellular calcium and H_2_O_2_ in the same root, the root was excited sequentially at 490 nm for GCAMP and 550 nm for Amplex Red with PE-4000 LEDs (CoolLed). Exposition times were chosen between 200 and 600 ms. The rootchip comprises 4 valves, allowing for imaging at best 4 positions of compression for 2 different roots (Supplementary Fig. 1). Time-lapse recordings were acquired either for one position only or with a multi-stage position setting, allowing for imaging the same root at different positions, under the valve or out of the valve. For the short and long compression with AmplexRed only, acquisition frequency was 1 frame every 5 to 20 s. For the kinetics experiments with H_2_O_2_ and calcium observed in the same root and the inhibition experiments, the acquisition frequency was 1 frame every 2 to 6 s. The exact time of acquisition for each frame was extracted from the image metadata.

Confocal cross-sections of the root were acquired using a Leica SP8X inverted confocal microscope equipped with a piezo stage (Super Z-galvo stage) allowing rapid z movement, a white light laser (WLL2, 470–670 nm) and hybrid detectors (GaAsP, Hamamatsu). Calcium and H_2_O_2_ were imaged in the same root with a 20X objective in water immersion (PL APO CORR CS2 multi-immersion, NA 0.7, Leica) and sequential acquisition, with λ_exc_ = 488 nm and λ_em_ = 500-530 nm for GCAMP, λ_exc_ = 550 nm and λ_em_ = 560-620 nm for Amplex Red, using 10%-20% laser power. For some roots, GCAMP or Amplex Red were imaged alone with a line average of 2 to keep a fast acquisition and reduce the signal to noise ratio. Roots expressing the ratiometric calcium probe RGECO-mTurquoise were observed with the same objective and the following sequential configuration: λ_exc_ = 570 nm/ λ_em_ = 575-650 nm for RGECO with gating (selection of photons depending on their lifetime) from 0.3 to 8 ns to avoid autofluorescence; λ_exc_ = 470 nm/ λ_em_ = 475-530 nm for mTurquoise with gating from 0.5 to 6 ns. Acquisition frequency was 1 frame every 2.78 s or 5.54 s, with or without bidirectional X setting.

### Image processing and data analysis

For wide-field images, the mean intensity of GCAMP and Amplex Red fluorescence was measured on manually defined ROI (region of interest) around the root using ImageJ (Fig. 1B). The same ROI was used for all frames of the same position, and the exact acquisition time of each frame was extracted from the image metadata. Confocal cross-sections with non-ratiometric probes were also directly analysed with ImageJ: ROIs for each root zone (epidermis, cortex and root center) were traced manually, and ROIs were traced again to track the same cells and tissues at each step of the deformation induced by pressure application (Fig. 1C). Mean intensity of fluorescence was measured on these ROIs. For ratiometric probe images, the ratio between RGECO (570 nm) and mTurquoise (470nm) was calculated with Python, using *skimage* and *numpy* modules and selecting ratio values between 0 and 10 to remove aberrant values outside of the root. The mean ratio for each root zone was then measured on manually defined ROIs on ImageJ.

From the mean ratios and mean intensity values for each frame and each position observed, 3 steps of analysis were performed in Python (pandas, numpy seaborn, matplotlib, pingouin and scipy.stats modules). The first step was the visualization of mean intensity values along the time for each acquisition. To compare responses in roots or root tissues with different basal intensity of the probes, a baseline value was defined as the mean of the values for 20 timepoints before the onset of pressure and the ratio between the mean intensity and the baseline was plotted over time for each replicate. The normalized intensity shown in the figures corresponds to this ratio, and the mean normalized intensity was calculated for different replicates with a confidence interval of 95 %. As the exact time after pressure start is not the same between images from different acquisitions, the normalized intensity values have been averaged on a time window of 20 seconds for the results shown in Figure 2 and 4 seconds for the rest of the results. Mean normalized intensity curves thus exhibit a slightly less precise time resolution compared to individual normalized intensity curves. For the second step, the amplitudes of H_2_O_2_ and calcium responses were defined as the difference between the value at 1 minute after pressure start and the minimum value right before pressure start. Amplitudes were calculated for the H_2_O_2_ wave and calcium slow wave as our time resolution was sufficient to get reliable values. As a third step of analysis, the slope of the mean intensity along the time curve was calculated. The time for which the slope becomes positive is considered the onset time of the response. The time for which the slope is maximal was also determined. In each slope curve, signal variations were identified as a positive slope followed by a slope decrease. Signal variations were then filtered with 3 criteria to determine if a peak was present and discriminate between the different phases: the time when the slope became positive, the peak duration and the end time must fit with the onset times, durations and end times obtained for the kinetic analysis. This allowed for quantification of the percentage of peaks or waves observed. Confidence intervals of the percentage of peaks were calculated with the Wilson method, which accounts for the small sample size. χ² independence test and pairwise χ² with Benjamini-Hochberg correction were used to test the hypothesis of the equality of the proportions. To compare the amplitude of the H_2_O_2_ response within the same root, repeated measurements ANOVA and paired T-test were used. When the distribution of values was non-parametric, for the H_2_O_2_ and calcium phases in the kinetics experiments, Friedman and pairwise Wilcoxon tests were used to compare kinetic parameters within the same root. To compare kinetic parameters after inhibition, as sample size varied between conditions, Welch ANOVA was used with post hoc Games-Howell test. Mann-Whitney U test was used for comparison between WT and mutants amplitudes. All boxplots show the median (middle bar), lower and upper quartiles (lower and upper bars of the box) and the whiskers represent the range of values.

## Acknowledgements

The authors would like to thank Sandrine Lécart, Romain Lebars, Amandine Dumazel and Valérie Nicolas from the Imagerie-Gif platform for their help (I2BC, Gif-sur-Yvette, France), advices and discussions. We thank Herman Höfte and Stefanie Afonso (IJPB, Versailles, France) who kindly provided *GCAMP* and *fer4 GCAMP* seeds, Jiri Friml and Yvan Kulich who kindly provided *rbohc/f* seeds (ISTA, Vienna, Austria) and Ting Li for providing piezo1-2 seeds (CalTech, Pasadena, USA). We thankAtef Asnacios (MSC, Paris, France) and Yvan Kulich for fruitful discussions.

## Funding

P. V. was funded by the “programme blanc” of the Graduate School BIOSPHERA, Université Paris-Saclay. This work benefited from Imagerie-Gif core facility supported by l’Agence Nationale de la Recherche (ANR-11-EQPX-0029/Morphoscope, ANR-10-INBS04/FranceBioImaging; ANR-11-IDEX-0003-02/Saclay Plant Sciences).

## Author contributions

P.V., S.T., J-M.F., P.D-S. and V.A. conceptualized the study and designed the experiments. P.V. and V.A. performed the experiments. P.V. conducted statistical and computational data analysis. P.V., S.T., J-M.F. and P.D-S. wrote the original draft. All authors edited the manuscript and approved the final manuscript version.

## Notes

### Competing Interest Statement

The authors have declared no competing interest.

### Summary of Updates

We performed experiments to ensure that the changes in the Amplex red signal we measure reflect H2O2 accumulation in response to root compression and that it originates from NADPH oxidase activity. We also tested mutants affecting mechanoperception and found that FERONIA is involved in the responses to lateral root compression.

## References

Alonso Baez L, Bjørkøy A, Saffioti F, Morghen S, Amanda D, Tichá M, Besten M, Ivanova A, Sprakel J, Stokke BT, et al (2026) The mechanical properties of Arabidopsis thaliana roots adapt dynamically during development and to stress. Science Advances 12: eaeb0032

Amador-Vargas S, Dominguez M, León G, Maldonado B, Murillo J, Vides GL (2014) Leaf-folding response of a sensitive plant shows context-dependent behavioral plasticity. Plant Ecol 215: 1445–1454

Audemar V, Guerringue Y, Frederick J, Vinet P, Melogno I, Babataheri A, Legué V, Thomine S, Frachisse J-M (2023) Straining the root on and off triggers local calcium signalling. Proceedings of the Royal Society B: Biological Sciences 290: 20231462

Basu D, Haswell ES (2020) The Mechanosensitive Ion Channel MSL10 Potentiates Responses to Cell Swelling in Arabidopsis Seedlings. Current Biology 30: 2716–2728.e6

Braam J, Davis RW (1990) Rain-, wind-, and touch-induced expression of calmodulin and calmodulin-related genes in Arabidopsis. Cell 60: 357–364

Demidchik V, Shabala SN, Coutts KB, Tester MA, Davies JM (2003) Free oxygen radicals regulate plasma membrane Ca2+- and K+-permeable channels in plant root cells. J Cell Sci 116: 81–88

Dong Q, Zhang Z, Liu Y, Tao L-Z, Liu H (2019) FERONIA regulates auxin-mediated lateral root development and primary root gravitropism. FEBS Letters 593: 97–106

Duan Q, Kita D, Li C, Cheung AY, Wu H-M (2010) FERONIA receptor-like kinase regulates RHO GTPase signaling of root hair development. Proceedings of the National Academy of Sciences 107: 17821–17826

Ermakov YA, Kamaraju K, Sengupta K, Sukharev S (2010) Gadolinium Ions Block Mechanosensitive Channels by Altering the Packing and Lateral Pressure of Anionic Lipids. Biophysical Journal 98: 1018–1027

Evans MJ, Choi W-G, Gilroy S, Morris RJ (2016) A ROS-Assisted Calcium Wave Dependent on the AtRBOHD NADPH Oxidase and TPC1 Cation Channel Propagates the Systemic Response to Salt Stress. Plant Physiol 171: 1771–1784

Fichman Y, Zandalinas SI, Peck S, Luan S, Mittler R (2022) HPCA1 is required for systemic reactive oxygen species and calcium cell-to-cell signaling and plant acclimation to stress. Plant Cell 34: 4453–4471

Fino LMD, Anjam MS, Besten M, Mentzelopoulou A, Papadakis V, Zahid N, Baez LA, Trozzi N, Majda M, Ma X, et al (2025) Cellular damage triggers mechano-chemical control of cell wall dynamics and patterned cell divisions in plant healing. Developmental Cell 60: 1411–1422.e6

Foreman J, Demidchik V, Bothwell JHF, Mylona P, Miedema H, Torres MA, Linstead P, Costa S, Brownlee C, Jones JDG, et al (2003) Reactive oxygen species produced by NADPH oxidase regulate plant cell growth. Nature 422: 442–446

Frachisse J-M, Thomine S, Allain J-M (2020) Calcium and plasma membrane force-gated ion channels behind development. Current Opinion in Plant Biology 53: 57–64

Gao D, Knight MR, Trewavas AJ, Sattelmacher B, Plieth C (2004) Self-Reporting Arabidopsis Expressing pH and [Ca2+] Indicators Unveil Ion Dynamics in the Cytoplasm and in the Apoplast under Abiotic Stress. Plant Physiol 134: 898–908

Guerringue Y, Thomine S, Allain J-M, Frachisse J-M (2026) The Rapid Mechanically Activated channel transduces increases in plasma membrane tension into transient calcium influx. New Phytologist 251: 276–287

Hartmann FP, Tinturier E, Julien J-L, Leblanc-Fournier N (2021) Between Stress and Response: Function and Localization of Mechanosensitive Ca2+ Channels in Herbaceous and Perennial Plants. International Journal of Molecular Sciences 22: 11043

Howell AH, Völkner C, McGreevy P, Jensen KH, Waadt R, Gilroy S, Kunz H-H, Peters WS, Knoblauch M (2023) Pavement cells distinguish touch from letting go. Nat Plants 1–6

Kolb E, Hartmann C, Genet P (2012) Radial force development during root growth measured by photoelasticity. Plant Soil 360: 19–35

Krogman W, Sparks JA, Blancaflor EB (2020) Cell Type-Specific Imaging of Calcium Signaling in Arabidopsis thaliana Seedling Roots Using GCaMP3. International Journal of Molecular Sciences 21: 6385

Kulich I, Vladimirtsev D, Randuch M, Gao S, Citterico M, Konrad KR, Nagel G, Wrzaczek M, Cascaro L, Vinet P, et al (2026) Calcium-triggered apoplastic ROS bursts balance gravity and mechanical signals for soil navigation. Science 392: 296–300

Lu B, Wang S, Feng H, Wang J, Zhang K, Li Y, Wu P, Zhang M, Xia Y, Peng C, et al (2024) FERONIA-mediated TIR1/AFB2 oxidation stimulates auxin signaling in Arabidopsis. Molecular Plant 17: 772–787

Malivert A, Hamant O (2023) Why is FERONIA pleiotropic? Nat Plants 9: 1018–1025

Marcec MJ, Tanaka K (2022) Crosstalk between Calcium and ROS Signaling during Flg22-Triggered Immune Response in Arabidopsis Leaves. Plants 11: 14

Martin L, Leblanc-Fournier N, Julien J-L, Moulia B, Coutand C (2010) Acclimation kinetics of physiological and molecular responses of plants to multiple mechanical loadings. J Exp Bot 61: 2403–2412

Mittler R, Zandalinas SI, Fichman Y, Van Breusegem F (2022) Reactive oxygen species signalling in plant stress responses. Nat Rev Mol Cell Biol 23: 663–679

Monshausen GB, Bibikova TN, Weisenseel MH, Gilroy S (2009) Ca2+ Regulates Reactive Oxygen Species Production and pH during Mechanosensing in Arabidopsis Roots. The Plant Cell 21: 2341–2356

Morré DJ (2002) Preferential Inhibition of the Plasma Membrane NADH Oxidase (NOX) Activity by Diphenyleneiodonium Chloride with NADPH as Donor. Antioxidants & Redox Signaling 4: 207–212

Ogasawara Y, Kaya H, Hiraoka G, Yumoto F, Kimura S, Kadota Y, Hishinuma H, Senzaki E, Yamagoe S, Nagata K, et al (2008) Synergistic Activation of the Arabidopsis NADPH Oxidase AtrbohD by Ca2+ and Phosphorylation *. Journal of Biological Chemistry 283: 8885–8892

Pei S, Tao Q, Li W, Qi G, Wang B, Wang Y, Dai S, Shen Q, Wang X, Wu X, et al (2024) Osmosensor-mediated control of Ca2+ spiking in pollen germination. Nature. doi: 10.1038/s41586-024-07445-6

Pei Z-M, Murata Y, Benning G, Thomine S, Klüsener B, Allen GJ, Grill E, Schroeder JI (2000) Calcium channels activated by hydrogen peroxide mediate abscisic acid signalling in guard cells. Nature 406: 731–734

Pomiès L, Decourteix M, Franchel J, Moulia B, Leblanc-Fournier N (2017) Poplar stem transcriptome is massively remodelled in response to single or repeated mechanical stimuli. BMC Genomics 18: 300

Ranf S, Eschen-Lippold L, Pecher P, Lee J, Scheel D (2011) Interplay between calcium signalling and early signalling elements during defence responses to microbe- or damage-associated molecular patterns. The Plant Journal 68: 100–113

Ravi B, Foyer CH, Pandey GK (2023) The integration of reactive oxygen species (ROS) and calcium signalling in abiotic stress responses. Plant, Cell & Environment 46: 1985–2006

Reis J, Massari M, Marchese S, Ceccon M, Aalbers FS, Corana F, Valente S, Mai A, Magnani F, Mattevi A (2020) A closer look into NADPH oxidase inhibitors: Validation and insight into their mechanism of action. Redox Biology 32: 101466

Richards SL, Laohavisit A, Mortimer JC, Shabala L, Swarbreck SM, Shabala S, Davies JM (2014) Annexin 1 regulates the H2O2-induced calcium signature in Arabidopsis thaliana roots. The Plant Journal 77: 136–145

Richter GL, Monshausen GB, Krol A, Gilroy S (2009) Mechanical Stimuli Modulate Lateral Root Organogenesis. Plant Physiology 151: 1855–1866

Shih H-W, Miller ND, Dai C, Spalding EP, Monshausen GB (2014) The Receptor-like Kinase FERONIA Is Required for Mechanical Signal Transduction in Arabidopsis Seedlings. Current Biology 24: 1887–1892

Smokvarska M, Bayle V, Maneta-Peyret L, Fouillen L, Poitout A, Dongois A, Fiche J-B, Gronnier J, Garcia J, Höfte H, et al (2023) The receptor kinase FERONIA regulates phosphatidylserine localization at the cell surface to modulate ROP signaling. Science Advances 9: eadd4791

Tan Y-Q, Yang Y, Zhang A, Fei C-F, Gu L-L, Sun S-J, Xu W, Wang L, Liu H, Wang Y-F (2020) Three CNGC Family Members, CNGC5, CNGC6, and CNGC9, Are Required for Constitutive Growth of Arabidopsis Root Hairs as Ca2+-Permeable Channels. Plant Comm. doi: 10.1016/j.xplc.2019.100001

Tichá M, Richter H, Ovečka M, Maghelli N, Hrbáčková M, Dvořák P, Šamaj J, Šamajová O (2020) Advanced Microscopy Reveals Complex Developmental and Subcellular Localization Patterns of ANNEXIN 1 in Arabidopsis. Front Plant Sci. doi: 10.3389/fpls.2020.01153

Tran D, Galletti R, Neumann ED, Dubois A, Sharif-Naeini R, Geitmann A, Frachisse J-M, Hamant O, Ingram GC (2017) A mechanosensitive Ca2+ channel activity is dependent on the developmental regulator DEK1. Nat Commun 8: 1009

Waadt R, Krebs M, Kudla J, Schumacher K (2017) Multiparameter imaging of calcium and abscisic acid and high-resolution quantitative calcium measurements using R-GECO1-mTurquoise in Arabidopsis. New Phytologist 216: 303–320

Yamanaka T, Nakagawa Y, Mori K, Nakano M, Imamura T, Kataoka H, Terashima A, Iida K, Kojima I, Katagiri T, et al (2010) MCA1 and MCA2 That Mediate Ca2+ Uptake Have Distinct and Overlapping Roles in Arabidopsis. Plant Physiology 152: 1284–1296

Yoshimura K, Iida K, Iida H (2021) MCAs in Arabidopsis are Ca2+-permeable mechanosensitive channels inherently sensitive to membrane tension. Nat Commun 12: 6074

Yu W-W, Chen Q-F, Liao K, Zhou D-M, Yang Y-C, He M, Yu L-J, Guo D-Y, Xiao S, Xie R-H, et al (2024) The calcium-dependent protein kinase CPK16 regulates hypoxia-induced ROS production by phosphorylating the NADPH oxidase RBOHD in Arabidopsis. The Plant Cell 36: 3451–3466

