## supplementary figures for "Spatiotemporal Analysis Reveals Mechanisms Controlling Reactive Oxygen Species and Calcium Interplay Following Root Compression"

### Supplementary Figures and Materials

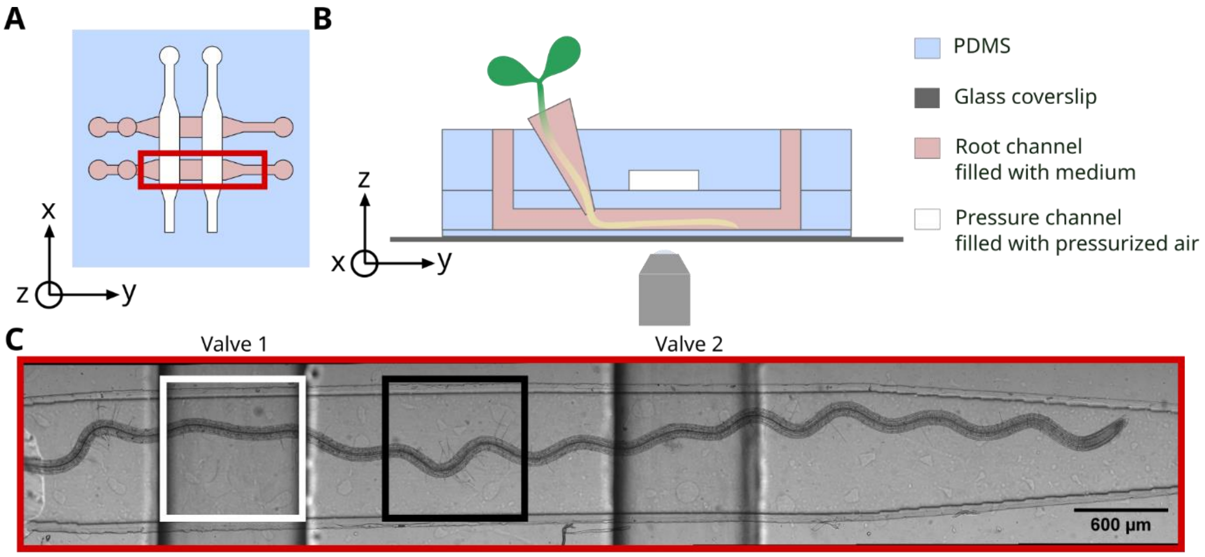


**Supplementary Figure 1: Microfluidic valve rootchip to observe root under lateral compression**

**(A)** Top view of a microfluidic rootchip with 2 root channels and 2 perpendicular pressure channels. Red rectangle corresponds to the imaged zone in **(C)**. **(B)** Side view of a microfluidic rootchip. **(C)** Bright-field image of a root inside the microfluidic rootchip, crossing 2 valves positions. White square corresponds to an “under the valve” imaging field. Black square corresponds to an “out of the valve” imaging field. Mosaic image obtained at 10X.

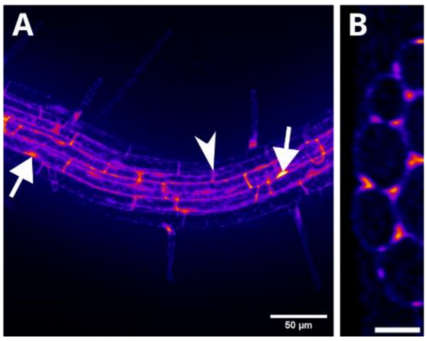


**Supplementary Figure 2: Amplex Red intracellular and extracellular localization**

**(A)** Z average projection on a xyz stack of 10 µM Amplex Red stained root (raw image, 20X), scale bar 50 µm. White arrows indicate nuclear and cytosolic localization of the probe. The white arrowhead shows a localization corresponding to the cell wall. **(B)** YZ plane of the deconvoluted xyz stack scale bar 20 µm. Color-coded intensity for (A and B).

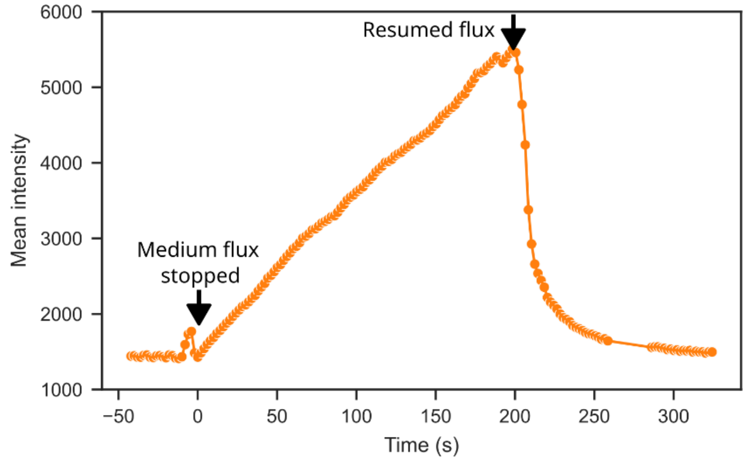


**Supplementary Figure 3 : Effect of interrupting and resuming Amplex Red flux on resorufin fluorescence intensity in the root**

Stopping the medium flux induces $H_2O_2$ accumulation in a root perfused with AmplexRed (black arrow). The syringe pump was started again after 200 seconds (second arrow), resulting in a rapid washout of the accumulated fluorescent resorufin by fresh medium.

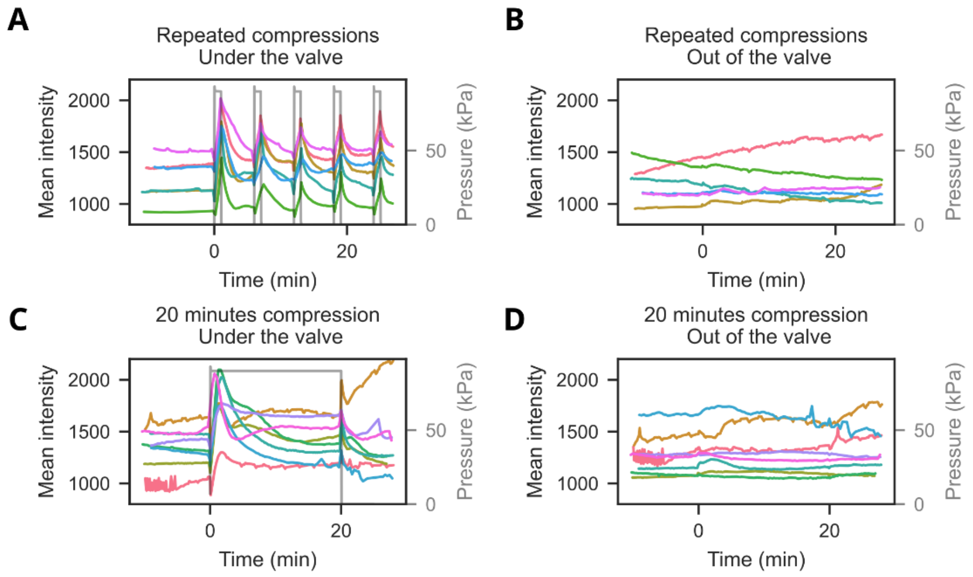


**Supplementary Figure 3: Mean intensity of $H_2O_2$ signal in response to repeated or long compressions**

Mean intensity of Amplex Red fluorescence signal for each root from 3 independent experiments, for positions under the pressure valve **(A)** **(C)** or out of the valve **(B)** **(D)** in response to 1 minute repeated compressions (A) or 20 minutes long compression (C). Same colors indicate same root replicate for **(A)** and **(B)** or **(C)** and **(D)**.

**Supplementary video 1: Kinetics of $H_2O_2$ and calcium responses in the root, Amplex Red and GCAMP channels**

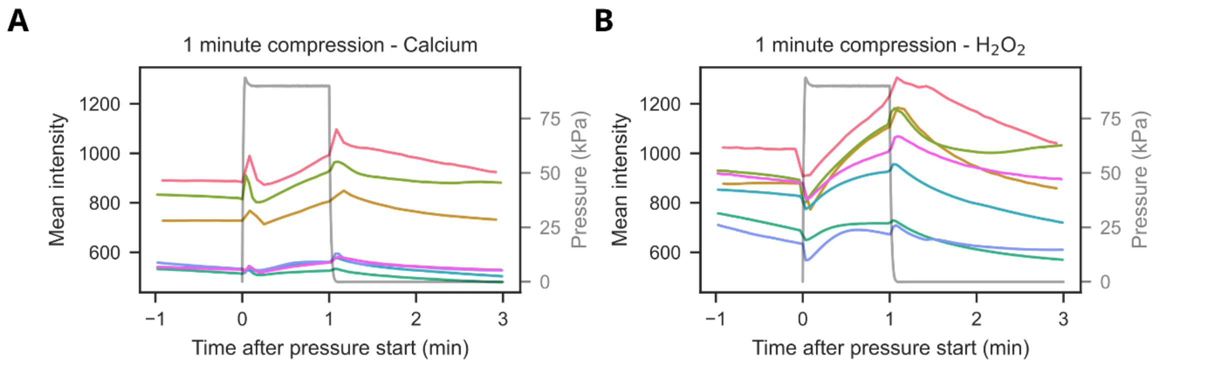


**Supplementary Figure 4: Mean intensity of Calcium and $H_2O_2$ probes in response to 1 minute compression**

Mean intensity of GCAMP calcium probe **(A)** and Amplex Red $H_2O_2$ probe **(B)** in response to 1 minute compression for each root from 3 independent experiments. Same colors indicate same root replicate.

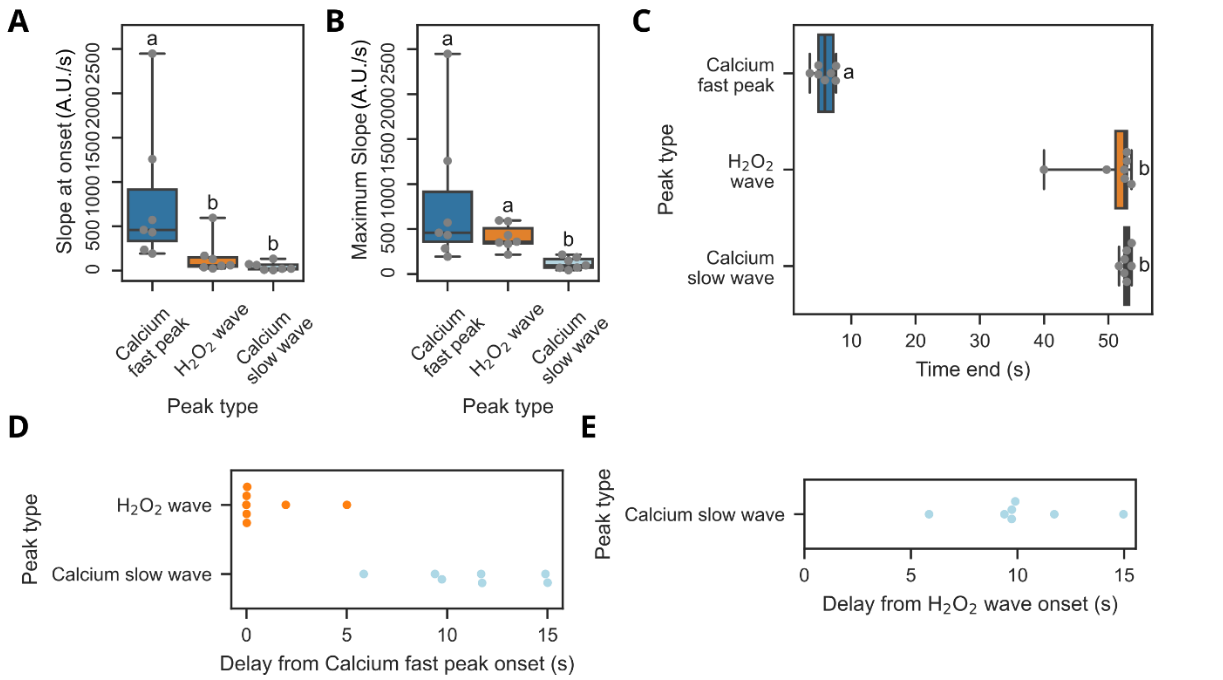


**Supplementary Figure 5: Kinetic parameters of calcium fast peak, ROS wave and calcium slow wave**

**(A)** Slope values at the beginning and **(B)** maximal slope values for the 3 kinetic phases. Different letters indicate significant difference, p-value < 0.05, Friedman test and pairwise Wilcoxon tests. N = 4 experiments, n = 7 positions on 7 different roots. **(C)** Time of signal decrease for the 3 phases. **(D)** Delay from calcium fast peak onset for the $H_2O_2$ wave onset and the calcium slow wave onset. **(E)** Delay after the $H_2O_2$ wave onset for the calcium slow wave onset.

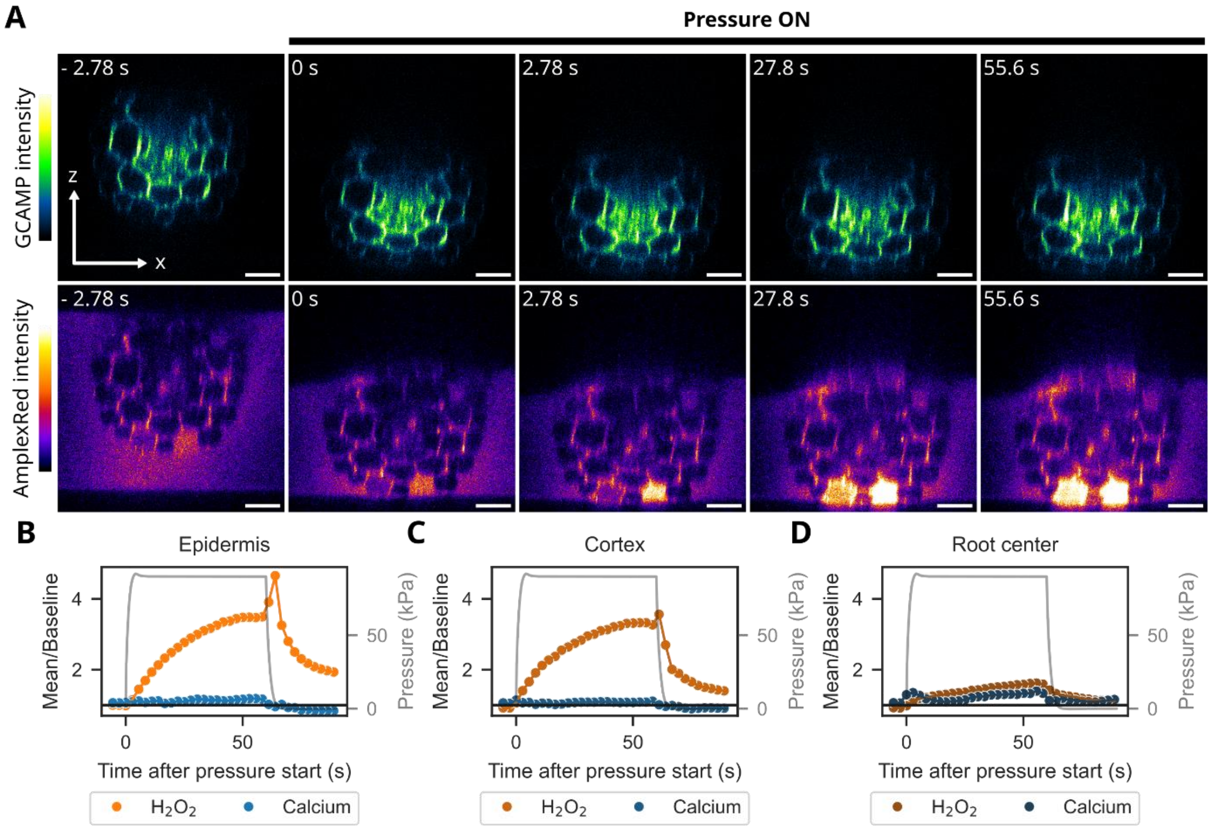


**Supplementary Figure 6: $H_2O_2$ and calcium responses localization in root n°2 (position 512)**

**(A)** Confocal cross sections of a root before and during a 1-minute compression, with color-coded fluorescence intensity of the GCAMP calcium probe (top) and Amplex Red ROS probe (bottom). The frames shown here are extracted from a time lapse acquisition, timestep of 2.78 s. Time origin is set at the onset of the pressure application. The root belongs to a different biological replicate than the one shown in Figure 4. **(B)**, **(C)**, **(D)** Ratio between the mean fluorescence intensity and the baseline intensity of $H_2O_2$ and calcium probes, in orange and blue, respectively. Baseline intensity is the mean of fluorescence intensity before pressure. This ratio is calculated on ROIs corresponding to the epidermis **(B)**, the cortex **(C)** and the root center **(D)** as illustrated in main Figure 1). The black line corresponds to a ratio of 1 in each ROI. The curves shown in B, C and D are the responses in the root shown in **(A)**.

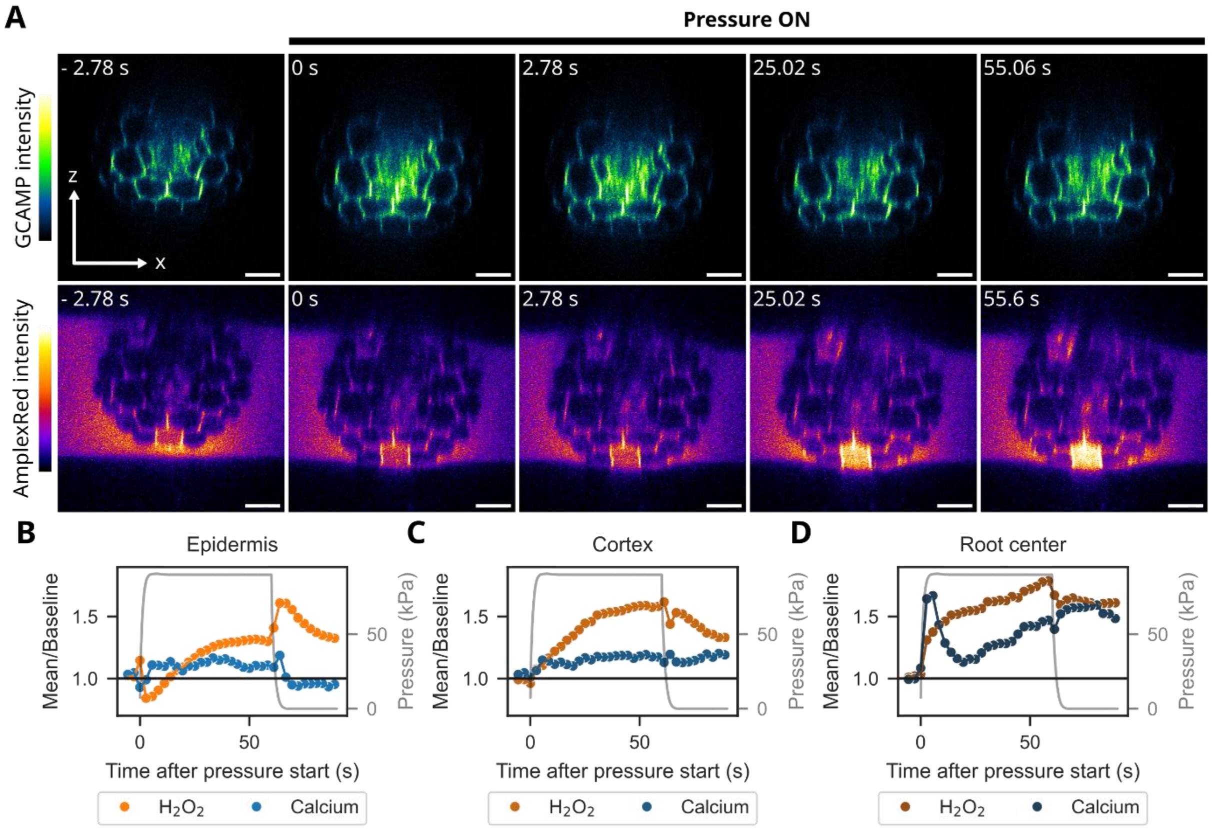


**Supplementary Figure 7: $H_2O_2$ and calcium responses localization in root n°3 (position 431)**

**(A)** Confocal cross sections of a root before and during a 1-minute compression, with color-coded fluorescence intensity of the GCAMP calcium probe and Amplex Red ROS probe in separated channels. The frames shown here are extracted from a timelapse acquisition, timestep of 2.78s. Time origin is set at the beginning of the pressure application. The root belongs to a different biological replicate than the ones shown in Figure 4 and S4. **(B)**, **(C)** and **(D)** Ratio between the mean fluorescence intensity and the baseline intensity of $H_2O_2$ and calcium probes, in orange and blue respectively. Baseline intensity is the mean of fluorescence intensity before pressure. This ratio is calculated on ROIs corresponding to the epidermis **(B)**, the cortex **(C)** and the root center **(D)**. The black line corresponds to a ratio of 1 in each ROI. The curves shown in B, C and D are the responses in the root shown in **(A)**.

**Supplementary video 2: Cross section of a root under compression, Amplex Red and GCAMP channels**

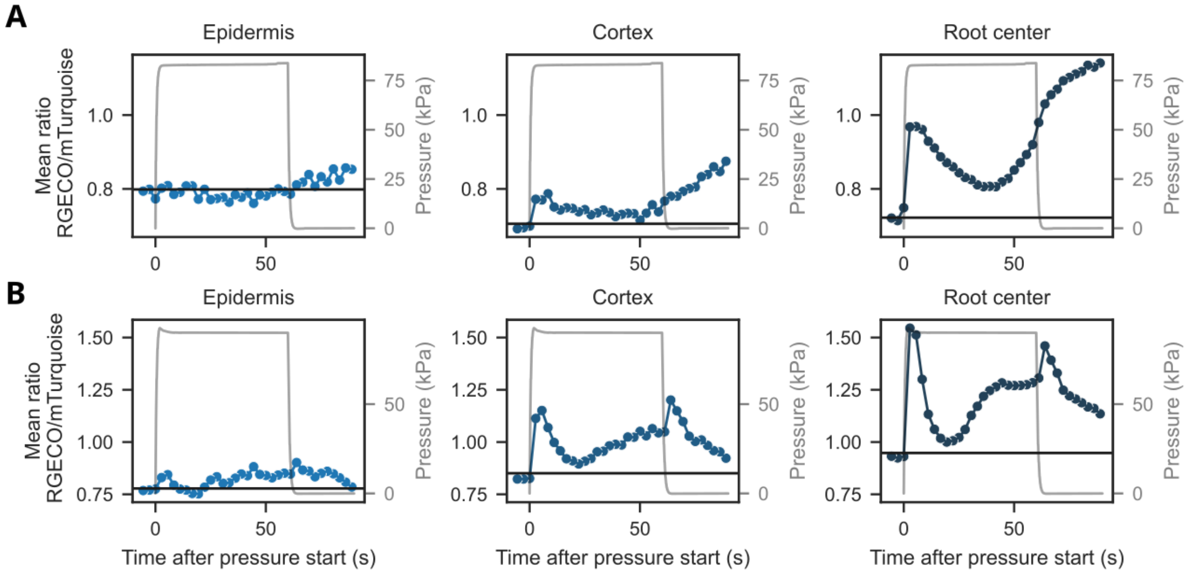


**Supplementary Figure 8: Calcium response localization in different root tissues measured with the ratiometric probe RGECO-mTurquoise**

**(A)** and **(B)** RGECO/mTurquoise mean fluorescence ratio in epidermis, cortex and root center of two different roots from the same experiment. Pressure values over time in grey. Black horizontal lines indicate the mean ratio baseline value.

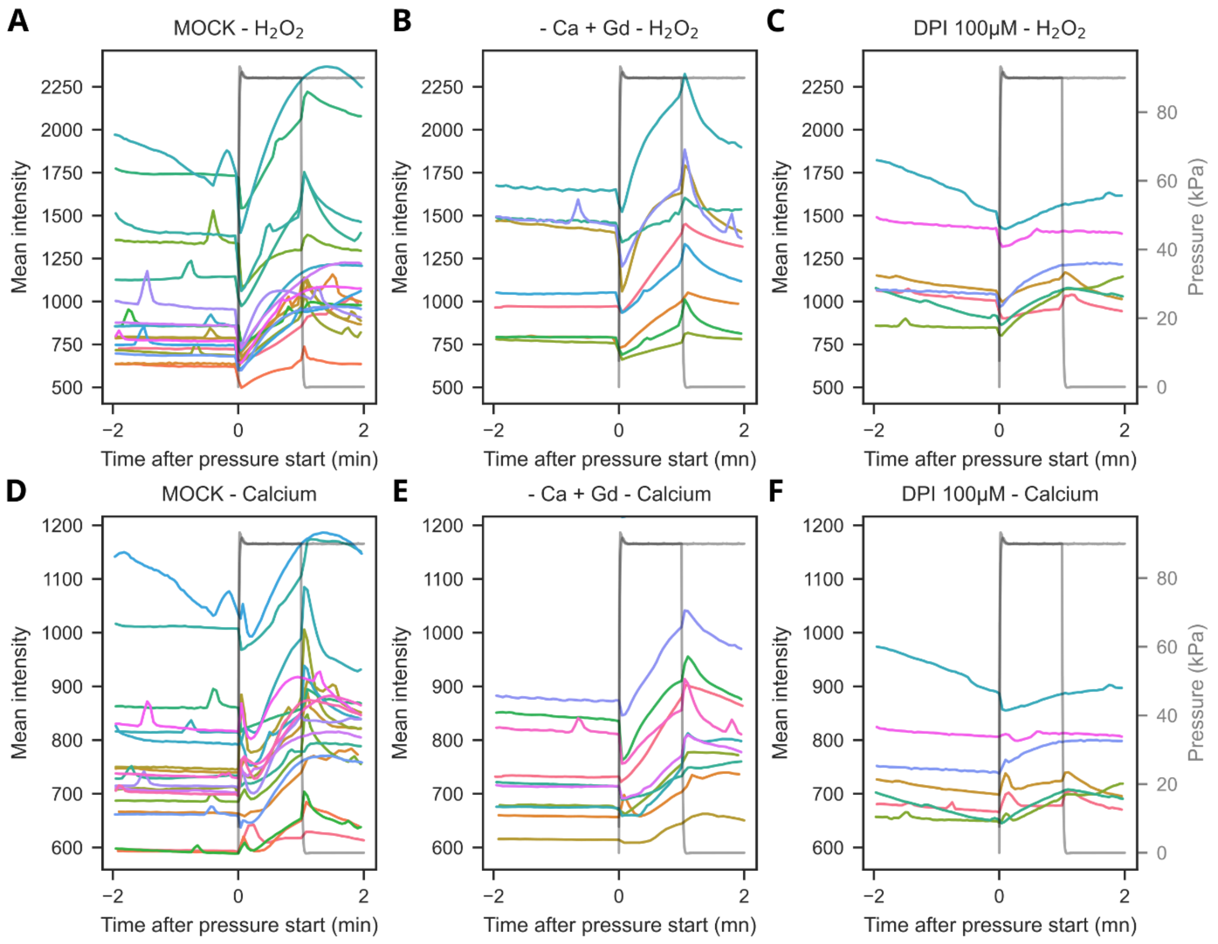


**Supplementary Figure 9: Mean intensity of calcium and $H_2O_2$ signals in response to compression in roots treated with inhibitors**

Mean intensity of Amplex Red **(A), (B), (C)** and GCAMP fluorescence **(D), (E), (F)** for each position of MOCK roots **(A), (D),** roots treated without calcium and 1 mM $Gd^{3+}$ **(B), (E)** or with 100 µM DPI **(C), (F)**. Each replicate from one condition corresponds to a different color. Based on these data from 3 independent experiments, the intensity was normalized to the baseline for each replicate and the mean of the normalized intensity curves is shown in Figure 5.

**Supplementary video 3: Calcium and $H_2O_2$ responses in a root treated with no calcium and 1 mM $Gd^{3+}$**

**Supplementary video 4: Calcium and $H_2O_2$ responses in a root treated with 100 µM DPI**

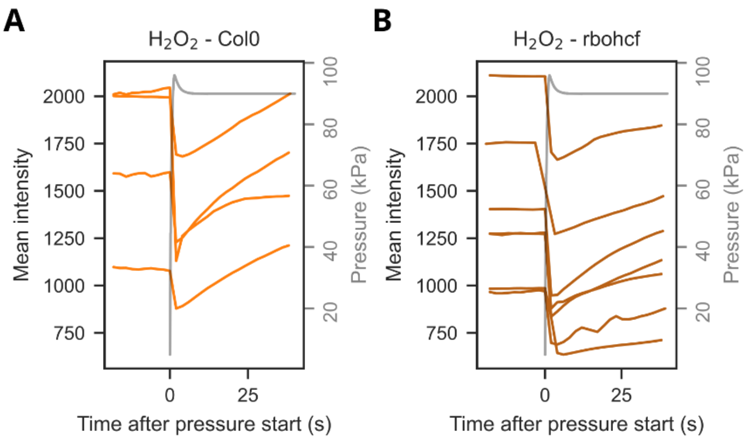


**Supplementary Figure 10: Mean intensity of $H_2O_2$ signal in response to compression in WT and *rbohcf* mutant**

Mean intensity of Amplex Red in Col0 **(A)** and *rbohc/f* roots **(B)**. Each curve corresponds to one position on one root. 4 roots were measured from 2 experiments for Col0, and 7 roots were measured from 3 experiments for *rbohc/f*.

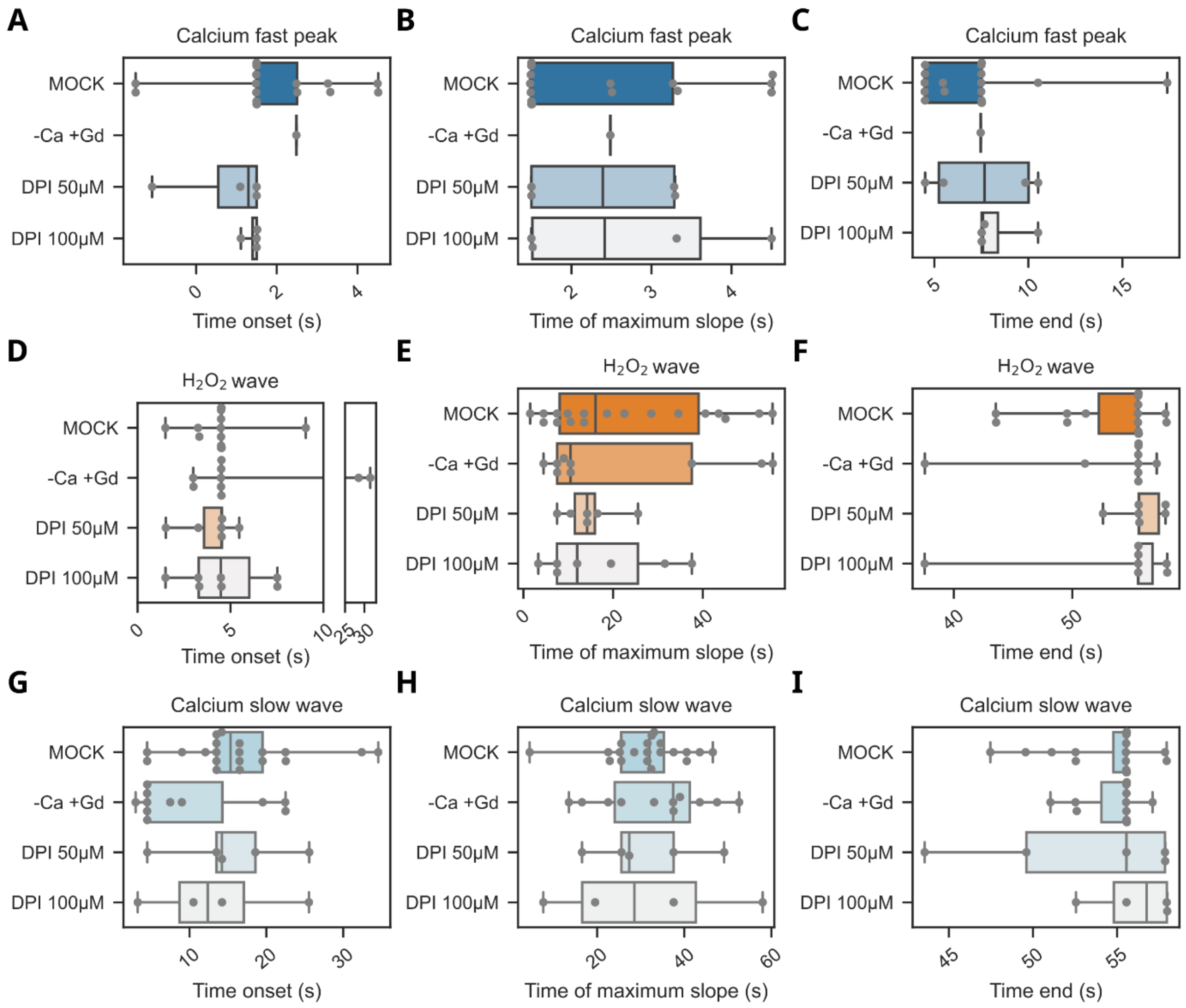


**Supplementary Figure 11: Time parameters of $H_2O_2$ and calcium response under Gd or DPI inhibition**

**(A), (D), (G)** Onset time, **(B), (E), (H)** time of maximum slope and **(C), (F), (I)** time of signal decrease of the calcium fast peak, $H_2O_2$ wave or calcium slow wave (respectively in **(A- C)**, **(D-F)** and **(G-I)**). These parameters were measured for a 1-minute compression of roots incubated in Hoagland medium (MOCK), Hoagland 0 calcium medium and 1 mM $Gd^{2+}$ (-Ca +Gd) or Hoagland medium and 100 µM DPI (100µM DPI). Calcium fast peak was detected in only one root out of 11 for -Ca +Gd condition. N.S., Welch ANOVA and Games-Howell pairwise test. N ≥ 3, n ≥ 4 positions on n ≥ 4 different roots for each treatment.

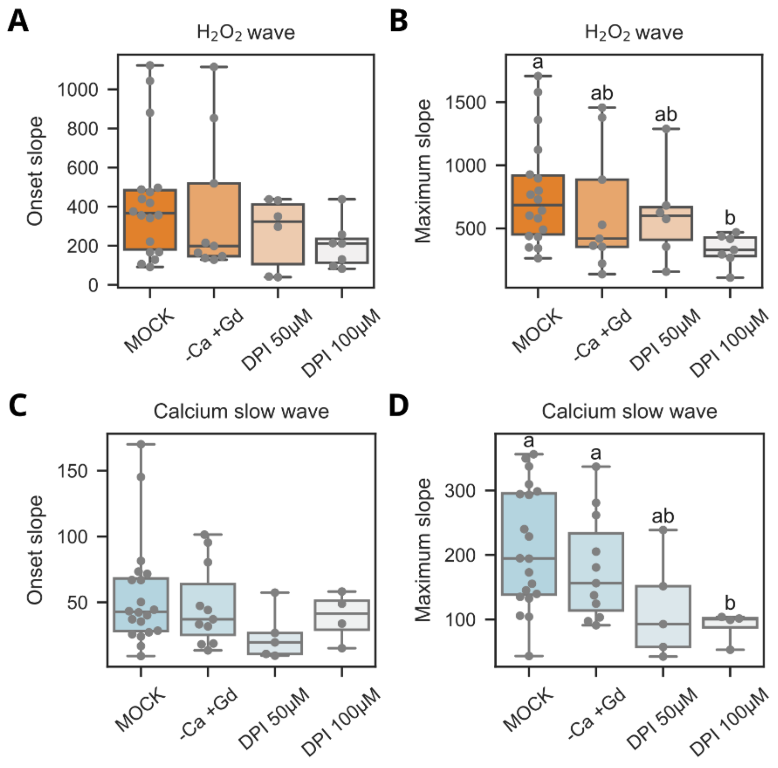


**Supplementary Figure 12: Slope parameters of $H_2O_2$ and calcium response under Gd or DPI inhibition**

**(A), (C)** Onset slope and **(B), (D)** maximum slope of the $H_2O_2$ wave or calcium slow wave (respectively in **(A) and (B)**, **(C) and (D)**). These parameters were measured for a 1-minute compression of roots incubated in Hoagland medium (MOCK), Hoagland 0 calcium medium and 1 mM $Gd^{2+}$ (-Ca +Gd) or Hoagland medium and 100 µM DPI (100µM DPI). N.S., Welch ANOVA and Games-Howell pairwise test. N ≥ 3, n ≥ 4 positions on n ≥ 4 different roots for each treatment.

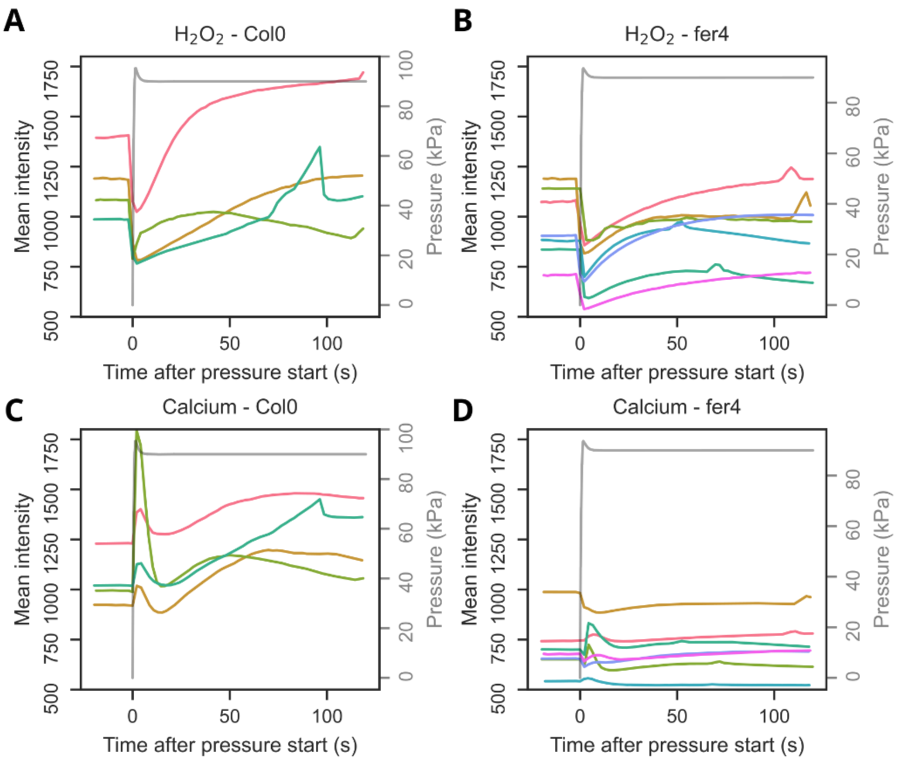


**Supplementary Figure 13: Mean intensity of calcium and $H_2O_2$ probes in response to compression in WT and *fer4* mutant roots**

Mean intensity of Amplex Red **(A), (B)** and GCAMP **(C), (D)** for each position of Col0 roots **(A), (C),** or *fer4* roots **(B), (D)**. Each replicate from one condition corresponds to a different color. Based on these data from 2 independent experiments for Col0 and 3 independent experiments for *fer4*, the intensity was normalized by the baseline for each replicate and the average of the normalized intensity curves is shown in Figure 6.

| Table 1: Kinetics parameters and amplitudes at 1 minute for the 3 response phases identified (mean ± standard deviation) | | | | | | | |
|---|---|---|---|---|---|---|---|
| **Experiment** | **Peak type** | **Time beginning (s)** | **Time maximum (s)** | **Time end (s)** | **Slope beginning** | **Maximum slope** | **Amplitude at 1 min** |
| Kinetics experiments (N = 4, n = 7 positions on 7 roots) | Calcium fast peak | 2.4 ± 0.94 | 2.7 ± 0.73 | 5.9 ± 1.5 | / | / | / |
| | ROS wave | 3.4 ± 2 | 7.3 ± 1.8 | 50.6 ± 4.8 | 153 ± 202 | 410 ± 139 | 186.4 ± 107.4 |
| | Calcium slow wave | 13.6 ± 6 | 29.3 ± 12.8 | 52.8 ± 0.7 | 47 ± 46 | 119 ± 65 | 64.4 ± 42.4 |
| Inhibition experiments MOCK (N = 10, n ≤ 18 positions on 16 roots ) | Calcium fast peak | 1.8 ± 1.6 | 2.4 ± 1.2 | 7.1 ± 3.1 | 490 ± 383 | 665 ± 643 | / |
| | ROS wave | 4.5 ± 1.4 | 23 ± 17.9 | 53.6 ± 4.3 | 426 ± 305 | 779 ± 423 | 372 ± 189 |
| | Calcium slow wave | 16.4 ± 7.5 | 31.2 ± 9 .2 | 54.6 ± 2.6 | 55 ± 40 | 212 ± 93 | 109 ± 47 |
| Inhibition experiments 0mM $Ca^{2+}$ + 1 mM $Gd^{3+}$ (N = 4, n ≤ 9 positions on 7 roots) | Calcium fast peak | / | / | / | / | / | / |
| | ROS wave | 9.8 ± 11.5 | 21.8 ± 21 | 53.2 ± 6.1 | 386 ± 365 | 638 ± 490 | 377 ± 201 |
| | Calcium slow wave | 9.7 ± 7.8 | 33.5 ± 12.6 | 54.8 ± 1.8 | 47 ± 31 | 180 ± 83 | 97 ± 45 |
| Inhibition experiments DPI 50µM (N = 3, n ≤ 5 positions on 5 roots) | Calcium fast peak | 0.8 ± 1.2 | 2.4 ± 1 | 7.6 ± 3 | / | / | / |
| | ROS wave | 4 ± 1.4 | 14.8 ± 8.2 | 55.8 ± 1.9 | 266 ± 282 | 613 ± 384 | 343 ± 210 |
| | Calcium slow wave | 15.3 ± 7.7 | 31.2 ± 12.5 | 52.9 ± 6.2 | 25 ± 20 | 117 ± 80 | 71 ± 37 |
| Inhibition experiments DPI 100µM (N = 4, n ≤ 6 positions on 6 roots) | Calcium fast peak | 1.4 ± 0.2 | 2.7 ± 1.5 | 8.3 ± 1.5 | / | / | / |
| | ROS wave | 4.6 ± 2.2 | 17 ± 13.1 | 53.7 ± 7.2 | 204 ± 122 | 332 ± 124 | 151 ± 96 |
| | Calcium slow wave | 13.4 ± 9.3 | 30.7 ± 22 | 56 ± 2.6 | 39 ± 19 | 89 ± 24 | 35 ± 19 |
